# Differential expression of single-cell RNA-seq data using Tweedie models

**DOI:** 10.1101/2021.03.28.437378

**Authors:** Himel Mallick, Suvo Chatterjee, Shrabanti Chowdhury, Saptarshi Chatterjee, Ali Rahnavard, Stephanie C. Hicks

**Affiliations:** Biostatistics and Research Decision Sciences, Merck & Co., Inc., Rahway, New Jersey 07065, U.S.A. *email:*; Epidemiology Branch, Division of Intramural Population Health Research, *Eunice Kennedy Shriver* National Institute of Child Health and Human Development, National Institutes of Health, Bethesda, MD 20892, USA *email:*; Department of Genetics and Genomic Sciences and Icahn Institute for Data Science and Genomic Technology, Icahn School of Medicine at Mount Sinai, New York, NY, 10029, USA *email:*; Global Statistical Sciences, Eli Lilly & Company, Indianapolis, IN 46225, U.S.A. *email:*; Computational Biology Institute, Department of Biostatistics and Bioinformatics, Milken Institute School of Public Health, The George Washington University, Washington, DC 20052, U.S.A. *email:*; Department of Biostatistics, Johns Hopkins Bloomberg School of Public Health, Baltimore, MD, 21205, U.S.A. *email:*

**Keywords:** Differential expression, Exponential dispersion model, Generalized linear model, Single-cell RNA-sequencing, Tweedie, Zero-inflation

## Abstract

The performance of computational methods and software to identify differentially expressed genes in single-cell RNA-sequencing (scRNA-seq) has been shown to be influenced by several factors, including the choice of the normalization method used and the choice of the experimental platform (or library preparation protocol) to profile gene expression in individual cells. Currently, it is up to the practitioner to choose the most appropriate differential expression (DE) method out of over 100 DE tools available to date, each relying on their own assumptions to model scRNA-seq data. Here, we propose to use generalized linear models with the Tweedie distribution that can flexibly capture a large dynamic range of observed scRNA-seq data across experimental platforms induced by heavy tails, sparsity, or different count distributions to model the technological variability in scRNA-seq expression profiles. We also propose a zero-inflated Tweedie model that allows zero probability mass to exceed a traditional Tweedie distribution to model zero-inflated scRNA-seq data with excessive zero counts. Using both synthetic and published plate- and droplet-based scRNA-seq datasets, we performed a systematic benchmark evaluation of more than 10 representative DE methods and demonstrate that our method (Tweedieverse) outperforms the state-of-the-art DE approaches across experimental platforms in terms of statistical power and false discovery rate control. Our open-source software (R package) is available at https://github.com/himelmallick/Tweedieverse.

## 1 Introduction

Advances in single-cell technologies now enable researchers to measure gene expression in tens of thousands of individual cells using single-cell RNA-sequencing (scRNA-seq) protocols, facilitating the understanding of a transcriptional landscape in unprecedented detail. In contrast to bulk RNA-sequencing (RNA-seq), scRNA-seq has been demonstrated to have a higher fraction of zeros per sample (or in this case cell), which can drive the primary source of variation and influence downstream analyses (Hicks et al., 2018; Ding et al., 2020). To date, there are over 20 library preparation protocols available (Chen et al., 2019) including the popular 10x Genomics Chromium (Zheng et al., 2017) or Drop-seq (Macosko et al., 2015) that both use unique molecular identifiers (UMIs) to remove PCR duplicate sequencing reads (Islam et al., 2014; Grün et al., 2014). In contrast, there are also plate-based, fulllength protocols, such as SMART-Seq2 (Picelli et al., 2014) or Fluidigm C1 (Pollen et al., 2014) that directly analyze read counts without UMIs, which can be useful, for example, to identify differential isoforms or allele-specific expression. We refer to scRNA-seq data from these two broad categories of protocols as *UMI counts* and *read counts* (or *non-UMI counts*).

Recent work has demonstrated that data generated from different scRNA-seq experimental protocols can result in differences in the distribution of gene expression, for example, between UMI counts and read counts, including differential zero-inflation or mean-variance patterns across experimental platforms (Vieth et al., 2017; Townes et al., 2019; Hafemeister and Satija, 2019; Svensson, 2020; Cao et al., 2021). This work emerged, in part, because historically many bioinformatics tools and statistical methods have been broadly proposed to model this technological variation in downstream analyses such as (i) zero-adjusted or zero-undjusted continuous models (Paulson et al., 2013; Korthauer et al., 2016; Soneson and Robinson, 2018), (ii) two-part hurdle models (Finak et al., 2015; Sekula et al., 2019), and (iii) count-based models such as Poisson, negative binomial, or multinomial models with (or without) zero-inflation components (Risso et al., 2018; Alessandrì et al., 2019; Hie et al., 2020).

Specifically in the context of differential expression (DE), many early methods designed for non-UMI counts focused on the use of zero-inflated or hurdle models (Finak et al., 2015; Korthauer et al., 2016; Miao et al., 2018; Sekula et al., 2019). Several of these methods require a data transformation including the use of a pseudocount and log-transformation, but this has been recently shown to introduce false variation in downstream analyses (Townes et al., 2019; Hafemeister and Satija, 2019). Furthermore, hurdle models can have restrictive assumptions that fail to distinguish between structural zeros and sampling zeros (Hu et al., 2011), which can be quite detrimental when the assumptions are violated. More recent methods designed for UMI counts proposed count-based models (Hafemeister and Satija, 2019; Townes et al., 2019), such as the negative binomial model, which has been classically used to model bulk RNA-seq data (Robinson et al., 2010; Love et al., 2014). However, recent work has questioned the general applicability of the negative binomial model for broad sequencing count data (Hawinkel et al., 2020). Currently, it is up to the practitioner to choose the most appropriate DE method out of approximately 100 DE tools available as of early 2021 (Zappia et al., 2018), each relying on their own assumptions, data transformations, statistical models, and inferential frameworks.

To address this, we propose a generalized framework that can flexibly capture a large dynamic range of observed scRNA-seq data across experimental platforms induced by heavy tails, sparsity, or different count distributions to model the technological variability in real-world scRNA-seq expression profiles. Specifically, we propose to use generalized linear models with a Tweedie distribution, parameterized by the mean (*μ*), dispersion (*φ*), and index or power parameter (*p*) (Zhang, 2013; Tweedie, 1984; Jørgensen, 1987). Our motivating reason for this distribution is that the index parameter *p* can flexibly capture the extent to which expression profiles differ according to the observed mean-variance relationships, including a Poisson (*p* = 1) or Compound Poisson *p* ∈ (1, 2) distribution. As a matter of fact, we found significant differences in gene-specific estimates of p when we considered 15 publicly available scRNA-seq datasets across experimental protocols and control experiments (**Fig. 1)**. In addition, our framework also considers a zero-inflated Tweedie model that allows zero probability mass to exceed a traditional Tweedie distribution to model zero-inflated scRNA-seq data with excessive zero counts. One of the main advantages of our approach is that it is technology-agnostic and can be applied to cross-platform scRNA-seq expression profiles with discrete (UMI or read) counts or normalized continuous expression levels, providing a unified framework for analyzing diverse scRNA-seq datasets. To the best of our knowledge, no previous work systematically studied the Tweedie models presented in the paper, particularly in the context of differential expression analysis of scRNA-seq data.

**Figure 1.**
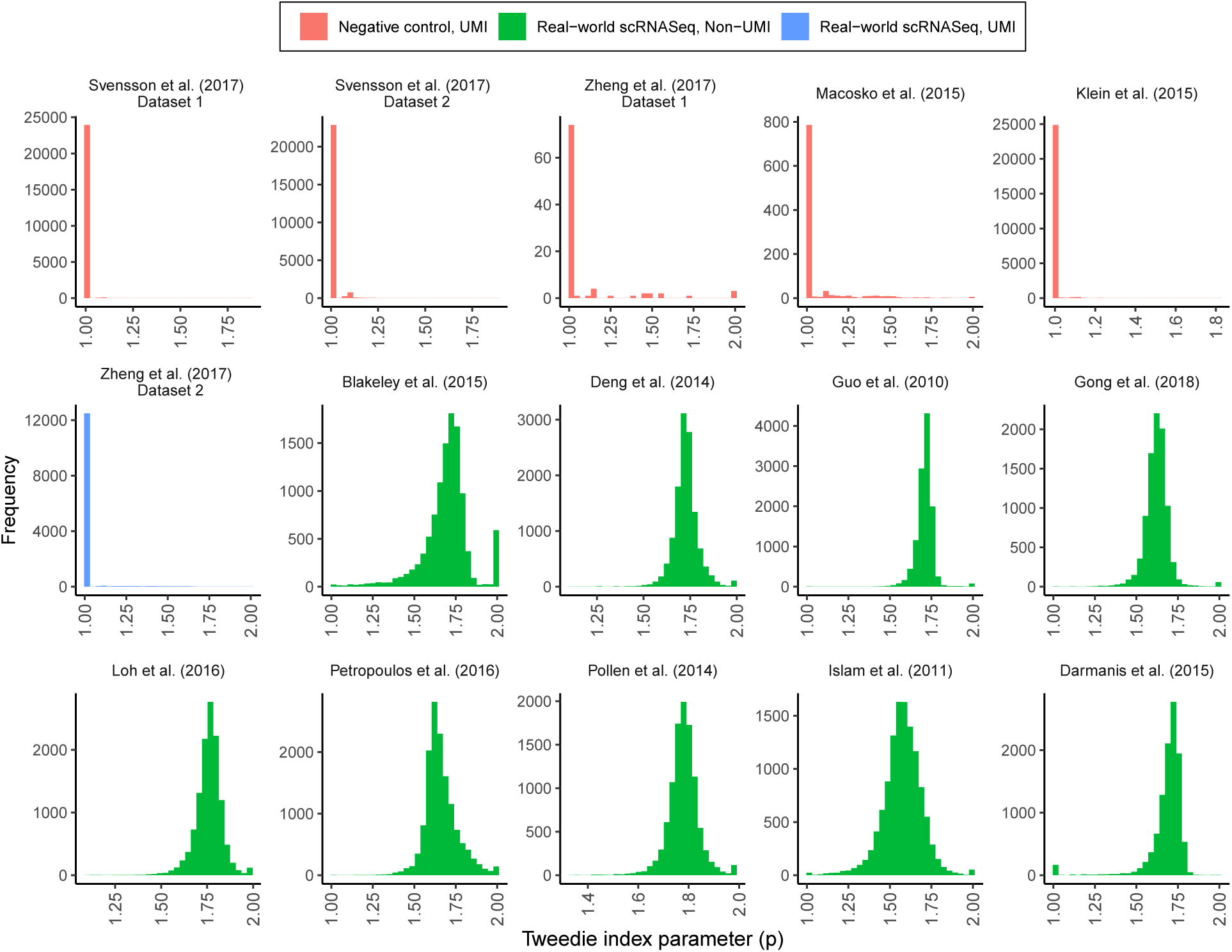
Histograms of gene-specific estimates of the Tweedie index parameter (*p*) vary across scRNA-seq experimental protocols and control datasets. Data from *N* = 15 publicly available scRNA-seq studies (full details of the datasets are described in **Web Table S1**) with (i) UMI counts or read counts and (ii) negative control experiments or real-world scRNA-seq experiments. A per-gene estimate of the Tweedie index parameter *p* was performed using maximum likelihood estimation. Using UMI count data, the majority of gene-wise estimates of *p* are at 1, which agrees with recent work suggesting that Poisson or multinomial models can be used to analyze these data. In contrast, using read count data, we observe a wide range of gene-specific estimates of *p*, suggesting that a flexible, generalized framework that can capture the technological variation is most appropriate for scRNA-seq DE analysis.

The rest of the manuscript is organized as follows. First, we define the three-parameter Tweedie family of distributions and introduce the resulting adaptive Tweedie models (both zero-inflated and non-zero-inflated) along with the algorithms for estimating the model coefficients (Section 2). Next, using both synthetic and published UMI and read count scRNA-seq datasets, we performed a systematic benchmark evaluation of more than 10 representative DE methods and demonstrate that our method (Tweedieverse) outperforms the state-of-the-art DE approaches across experimental platforms in terms of statistical power and false discovery rate control (Section 3). Finally, we conclude with a brief discussion of our results (Section 4). Our open-source software package is available at https://github.com/himelmallick/Tweedieverse.

## 2 Methodology

In this section, we first briefly describe the Tweedie distribution (Section 2.1) and the extension to a zero-inflated Tweedie distribution (Section 2.2). Next, we describe our proposed generalized model for differential expression using either distribution (Section 2.3) along with our testing strategy (Section 2.4). There are several advantages of our approach. First, by utilizing the library size (or scale factor) as an offset, we bypass the need for an *ad hoc* data transformation prior to DE testing. Second, we estimate a gene-specific index (power) parameter (*p*) that adapts to differences in mean-variance patterns within a given dataset, leading to substantial power gains in detecting DE genes. Third, our approach is technology-agnostic and can be applied to cross-platform scRNA-seq expression profiles, such as discrete counts (both UMI counts and read counts) as well as normalized continuous expression levels (in sharp contrast to existing methods that are either count-based or continuous but not both), providing a unified framework for analyzing diverse scRNA-seq datasets.

### 2.1 Tweedie distribution

Let *Y_gc_* denote the gene expression measurement of gene *g* (*g* = 1,…, *G*) in cell *c* (*c* = 1,…,*N*), where *Y_gc_* can be either UMI counts or read counts. Depending on whether the scRNA-seq data are UMI counts (previously shown to be non-zero-inflated) or read counts (previously shown to be zero-inflated), we model *Y_gc_* using either a Tweedie (*Tw*) distribution (i.e. *Y_gc_ ~ Tw*(*μ_gc_,ϕ_g_,p_g_*)) or a zero-inflated Tweedie (*ZITw*) distribution (i.e. *Y_gc_ ~ ZITw*(*μ_gc_,ϕ_g_,p_g_*)) respectively, where *μ_gc_* is the mean expression for gene *g* in cell *c* and *ϕ_g_* and *p_g_* denote the corresponding gene-specific dispersion and power parameters, satisfying *ϕ_g_* > 0 and *p_g_ ∈* (1,2). With a slight abuse of notation, we drop the subscripts *g* and *c* in the remainder of the manuscript.

An advantage of the Tweedie family is that it encompasses a number of well-known distributions for various choices of the power parameter such as the Normal distribution (*p* = 0), Poisson (*p* = 1), Gamma (*p* = 2), and Inverse Gaussian (*p* = 3) distributions, among others (Kurz, 2017). Our interest in using the Tweedie distribution is motivated by the fact that when 1 < *p* < 2, the Tweedie distribution reduces to a semi-continuous distribution with a point mass at zero followed by a right-skewed distribution with positive support. In the semi-continuous support range *p* ∈ (1,2), the Tweedie distribution is characterized by a mixture of Poisson and a compound sum of i.i.d. Gamma random variables, i.e.,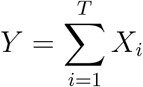 and *T ~ Pois*(*λ*), where 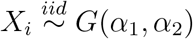 and *T* ⊥ *X_i_*, ∀*i*. Notably, for *p* ∈ (1,2), the Tweedie distribution is also called the Compound Poisson-Gamma distribution.

The positive probability mass at zero and a skewed distribution on the positive real line makes the Tweedie distribution an intuitive alternative for fitting both count and positive continuous data in the presence of exact zeros. This makes it particularly suitable for highly skewed, sparse UMI or read counts without the need to introduce zero-inflation or hurdle components, by the simple estimation of the index parameter. Moreover, by estimating a gene-specific index parameter, the Tweedie distribution flexibly captures a variable range of observed scRNA-seq expression profiles spanning a range of mean-variance relationships and sparsity patterns across platforms (**Fig. 1**). As detailed in the Supporting Information (**Web Appendix S1**), the re-parametrization of the Tweedie distribution as a member of the exponential dispersion model (EDM) family allows us to utilize the well-studied theory of generalized linear models (GLMs) with a Tweeedie distribution for the response.

### 2.2 Zero-inflated Tweedie distribution

One common feature of scRNA-seq read counts is the presence of excess zeros compared to what we expect under a particular count model causing many transcripts to go undetected for technical reasons, such as inefficient cDNA polymerization, amplification bias, or low sequencing depth (Van den Berge et al., 2018). In this case, the observed zeros in read count data can occur as a result of both biological and technical reasons. Therefore, a traditional Tweedie model is unsatisfactory in the presence of excess zeros. For zero-inflated scRNA-seq gene expression profiles, we propose to use a zero-inflated analogue of the Tweedie distribution. To our knowledge, this is the first study investigating the use of a zero-inflated Tweedie distribution and its estimation framework, both in the application of scRNA-seq DE analysis and otherwise.

Similar to other zero-inflated model formulations, a *ZITw* distribution assumes a mixture of Tweedie model component with probability (1-*q*) and an exact zero mass with probability *q*, 0 ≤ *q* ≤ 1:

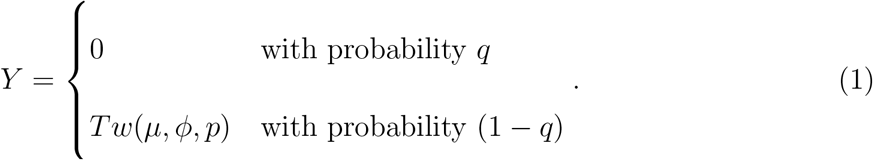

Under this specification, the probability of observing zeros is 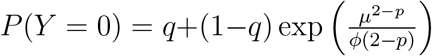, which is clearly greater than the probability of observing a zero count from the Tweedie distribution, indicating zero-inflation. Note that, the above-mentioned mixture model framework subsumes the traditional Tweedie model as a special case when no zero-inflation is present (i.e. *q* = 0).

### 2.3 Model description

To associate gene expression profiles with covariates using the Tweedie distribution, we parameterize the vector of per-gene mean count expression ***μ*** through the log link function as

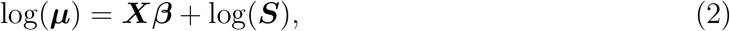

where ***S*** is the library size (or scale factor) included in the model as an offset, ***X*** is the design matrix, and ***β*** = (*β*_0_, *β*_1_,…, *β_k_*)’ is the vector of *k* regression coefficients, including the intercept. Of note, here we present a general case where the design matrix ***X*** (which is assumed to be of full rank) have *k* columns corresponding to *k* variables and hence ***β*** is a *k*-dimensional vector of regression coefficients. For the simple case of differential expression analysis, ***X*** is a two-column matrix consisting of an intercept and a binary variable indicator of group membership (e.g. healthy vs. control, biological cell types, etc.).

For assessing differential expression using the *ZITw* distribution, we additionally express the probability of zeros as a function of the predictors through a logit link and parameterize the mean vector through a log link function (Zhang, 2013) as before:

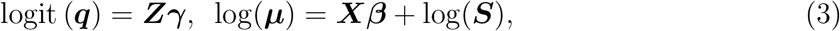

where ***q*** = (*q*_1_,…, *q_n_*) is the probability of observing excess zeros, ***Z*** and ***X*** are the design matrices corresponding to the two sub-components of the model, and ***γ*** and ***β*** are the vectors of regression coefficients for the zero-inflation and Tweedie components respectively. For simplicity, here we assume that the design matrices ***X*** and ***Z*** have the exact *k* columns corresponding to *k* covariates but in practice, they can be different.

### 2.4 Differential Expression Testing

To investigate differential expression for a given gene *g* (*g* = 1,…, *G*), we test the null hypothesis *H_0_*: *β_g_* = 0 against *H_1_*: *β_g_* ≠ 0 based on a univariate Tweedie model. We denote the estimate of *β_g_* by 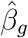, which is obtained by the profile likelihood approach (**Web Appendix S2**) when *y_i_ ~ Tw* or by the EM algorithm (**Web Appendix S3**) when *y_i_ ~ ZITw*, respectively. For the zero-inflated model, we specify an intercept-only design matrix for the zero subcomponent. For both models, we calculate the corresponding *p*-value based on the Wald’s test statistic, which, under the null hypothesis, follows an asymptotic χ^2^-distribution with 1 degree of freedom. We apply standard methods for controlling the False Discovery Rate (FDR) (described in detail in Section 3.1) and infer a gene to be differentially expressed between two experimental conditions if the FDR-adjusted *p*-value (*q*-value) falls below a pre-specified Type 1 error threshold (e.g. *α* = 0.05).

In addition to the gene-level coefficients, *p*-values, and *q*-values, we also report the estimated index parameter 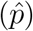, which is obtained as an automatic byproduct of our algorithm. The per-gene index parameter provides useful information about the distribution of the overall gene expression as it varies according to the observed sparsity and mean-variance pattern in the dataset (**Fig. 1**). As shown in Section 3, by optimizing the index parameter on a per-gene basis, our proposed Tweedie models are able to capture a wide range of distributions observed in multi-platform scRNA-seq expression profiles, resulting in a boost in statistical power while maintaining FDR across a variety of real and synthetic datasets. In what follows, we refer to our Tweedie model as Compound Poisson Linear Model (CPLM) and the zero-inflated Tweedie model as Zero-inflated Compound Poisson Linear Model (ZICP) due to their close connection to the Compound Poisson distribution and GLMs. We use the `glm()` and `optim()` functions of the *stats* base R package, and the *cplm* (Zhang, 2013), *statmod* (Giner and Smyth, 2016), and *tweedie* (Dunn and Smyth, 2005, 2008) R packages to carry out the above-mentioned computation. We have implemented our proposed methods in an R package called *Tweedieverse*. The software, documentation, and example datasets are made freely available at https://github.com/himelmallick/Tweedieverse.

## 3 Results

First, we present practical implementation details related to Tweedieverse (Section 3.1). Second, we present simulation studies that are designed to assess the performance of Tweedieverse in realistic synthetic data (in both UMI counts and read counts) compared to other state-of-the-art methods (Section 3.2). Finally, we apply Tweedieverse to a variety of real scRNA-seq datasets to identify biological discoveries that otherwise cannot be revealed by existing approaches (Section 3.3). Additional numerical results are presented in the Supporting Information.

### 3.1 Implementation details

Tweediverse expects either UMI or read counts that have been preprocessed using standard scRNA-seq cell- and gene-specific quality control metrics, normalization, and batch correction methods (as needed), described in Amezquita et al. (2019). These steps minimize the influence of unwanted technical and/or biological artifacts on the DE results. In particular, we first estimate cell-specific library sizes using the *scran* (Lun et al., 2016) R/Bioconductor package and incorporate them as an offset in the per-gene Tweedie models to account for differences in library sizes.

The *scran* package uses a deconvolution approach for normalization that initially partitions cells into pools, normalizes across cells within each pool, then uses the resulting system of linear equations to define cell library sizes. Additionally, it takes extra steps to counteract global assumptions about DE genes by estimating a library size term for each cell pool and utilizing these estimates to approximate a robust size factor term for individual cells. It must be emphasized that the choice of the normalization method can have a decided impact on the downstream analysis results and alternative normalization scale factors from other normalization methods can also be incorporated into our model as offset; however, we chose this method because of previous studies that have demonstrated superior performance over other tested normalization methods in the context of robust size factor estimation, batch effect correction, and differential expression (Luecken and Theis, 2019).

Unless otherwise stated, in the results presented in Sections 3.2–3.3, we apply the Benjamini-Yekutieli (Benjamini and Yekutieli, 2001) procedure to control the FDR (at the nominal *α* = 0.05 cutoff), which is a recommended procedure under arbitrary dependence of the tests. While the Benjamini-Hochberg adjustment (Benjamini and Hochberg, 1995) is commonly used to correct for multiple testing in scRNA-seq DE analysis, theoretical results only guarantee FDR control when the test statistics are independent or have a certain form of dependency termed positive regression under the null hypothesis. However, we and others observed that there is an appreciable amount of gene-gene correlations (also termed as ‘gene co-expression’) in scRNA-seq data - a phenomenon that is surprisingly consistent across real and synthetic datasets regardless of the data generation process or experimental platform (Assefa et al., 2020).

To substantiate this claim, we estimated the gene-gene correlations in a variety of real and simulated datasets. Following Assefa et al. (2020), we used the R package *WGCNA* (Langfelder and Horvath, 2008) to estimate the pairwise Pearson correlations (*r*) between the individual genes on the log-transformed counts per million (CPM) values (adding a small pseudo count of 1 to the zero observations before applying the log transformation). The distributions of the gene-to-gene correlations from the real and simulated scRNA-seq datasets reveal that there is a substantial amount of between-genes correlations exceeding Cohen’s (Cohen, 2013) recommended moderate effect size (i.e. *r* = 0.3), which we use to justify that the traditional BH adjustment is not appropriate for scRNA-seq per-gene DE testing (**Web Fig. S1**).

### 3.2 Simulation Studies

We benchmarked our methods using realistic synthetic datasets generated using the Splatter simulation framework (Zappia et al., 2017). Briefly, Splatter uses a parametric Poisson-gamma model to simulate scRNA-seq count data by estimating gene-specific parameters from a real-world template dataset. We used a variety of templates representing both read counts and UMI counts in order to mimic real-world scRNA-seq datasets. To this end, we used the Islam et al. (2011) dataset as a template for generating synthetic read counts, whereas the Klein et al. (2015) dataset was used as a template to generate synthetic UMI counts (**Web Table S1**). Within each of these settings, we further considered 8 core scenarios by varying (i) the zero-inflation mode (Yes/No), (ii) the number of cells (200, 500), and (iii) the number of genes (2000, 5000), each time summarizing performance over 100 simulation runs (**Web Fig. S2**). The location parameter for the differential expression factor (*de.facLoc*) was set to 1, which led to realistically varying log_2_-fold changes between (−3, 3) representing both modest (e.g. < 2-fold differences) and strong (e.g. 8-fold) effect sizes (**Web Fig. S3**). Values for the remaining parameters were either set to their default values or estimated from the respective template data. The probability of a gene being DE was set to 0.1, and cells had equal probabilities of being assigned to one of the two groups. Genes that were not expressed in at least 20% of the cells were removed from analysis.

We compared the performance of our proposed methods to a range published differential expression analysis tools. Specifically, we considered (i) four representative methods from the scRNA-seq literature: MAST (Finak et al., 2015), monocle (Trapnell et al., 2014), scREhurdle (Sekula et al., 2019) (NRE version), and DESingle (Miao et al., 2018); (ii) two methods designed for bulk RNA-seq differential expression: edgeR (Robinson et al., 2010) and DESeq2 (Love et al., 2014); (iii) one method specially geared towards metagenomics differential abundance analysis: metagenomeSeq (Paulson et al., 2013), and (iv) finally, a commonly used non-parametric method: Wilcoxon test. DE genes were determined at a FDR of 0.05 for each method.

We considered a variety of performance metrics including the F1 score (harmonic mean of precision and sensitivity), true positive rate (TPR) (proportion of detected DE genes among all true DE genes), and FDR (proportion of false positives among the declared significant findings), among others. In addition to the threshold-dependent performance metrics described above, we also considered threshold-agnostic evaluation metrics by varying the significance level between 0 and 1 to produce a series of mean TPRs and FDRs, which we subsequently use to obtain the receiver operating characteristic (ROC) curve as well the area under the ROC curve (AUC). We host our benchmarking code results in a repository that is aimed at simplifying future method evaluation and reanalysis of published results using new methods at https://github.com/himelmallick/BenchmarkSinglecell.

Our results revealed that both CPLM and ZICP consistently produced high F1 and AUC scores while controlling the FDR, with ZICP performing slightly better than CPLM in zero-inflated non-UMI counts (**Fig. 2**). All methods performed similarly in non-zero-inflated UMI counts, consistent with the recent observation that traditional non-zero-inflated count models are sufficient to model these data (Townes et al., 2019; Hafemeister and Satija, 2019). All methods maintained the Type I errors reasonably well (results not shown) and they all exhibited good, robust FDR control (with the exception of metagenomeSeq and monocole which are omitted due to their consistent FDR inflation across scenarios).

**Figure 2.**
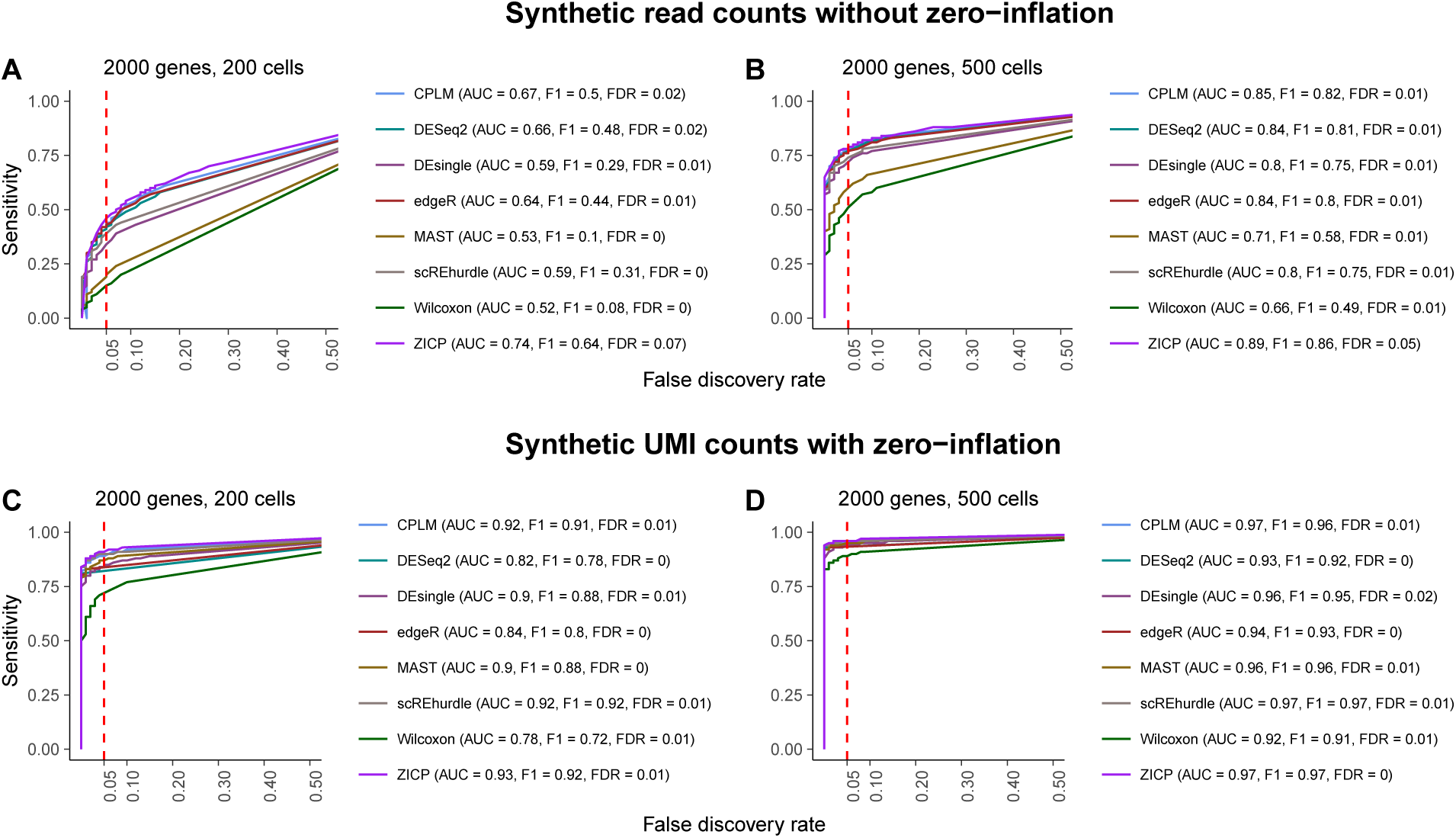
Performance results (Sensitivity) of Tweedieverse compared to existing DE methods using synthetic counts with and without zero-inflation across observed FDR. The performance of differential expression (DE) methods are compared using Splatter-simulated (**A, B**) synthetic read counts and (**C, D**) UMI counts depicting trade-offs between true positive rate (TPR or Sensitivity) (*y*-axis) and observed false discovery rate (FDR) (*x*-axis). The red dotted line represents *α*-level cutoff at *α* = 0.05. The area under the receiver operating characteristic (ROC) curves (AUC) and F1 scores are calculated at α = 0.05. Both CPLM and ZICP consistently resulted in the highest AUC values and F1 scores across read count data with a (**A**) smaller (*N* = 200) and (**B**) larger (*N* = 500) number of cells per group, and (**C**) UMI count data with a smaller (*N* = 200) number of cells. All methods performed similarly for (**D**) UMI count data with a larger (*N* = 500) number of cells. With the exception of metagenomeSeq and monocole (omitted due to their significant FDR inflation), nearly all methods controlled the FDR well below the imposed level. The red line parallel to the y-axis is the nominal threshold for FDR in multiple testing. Values are summarized averages over 100 iterations. The number of simulated genes and cells for each scenario are reported in the title of each panel.

When comparing our methodology to other methods for detecting DE genes, Tweedieverse remained one of the best-performing methods across various significance thresholds, a phenomenon that was consistent across all simulation scenarios (**Fig. 3**). Notably, the performance of the Tweedie models was found to be quite robust, which is generally unaffected by the dimensionality of the data (**Web Figs. S4-S5**) as well as the presence and absence of zero-inflation (**Web Figs. S6-S9**) in both UMI counts and read counts. Combined, these results further highlight the advantage of using a generalized model that adapts to the underlying characteristics of diverse single-cell expression profiles leading to substantial power gain in detecting DE genes.

**Figure 3.**
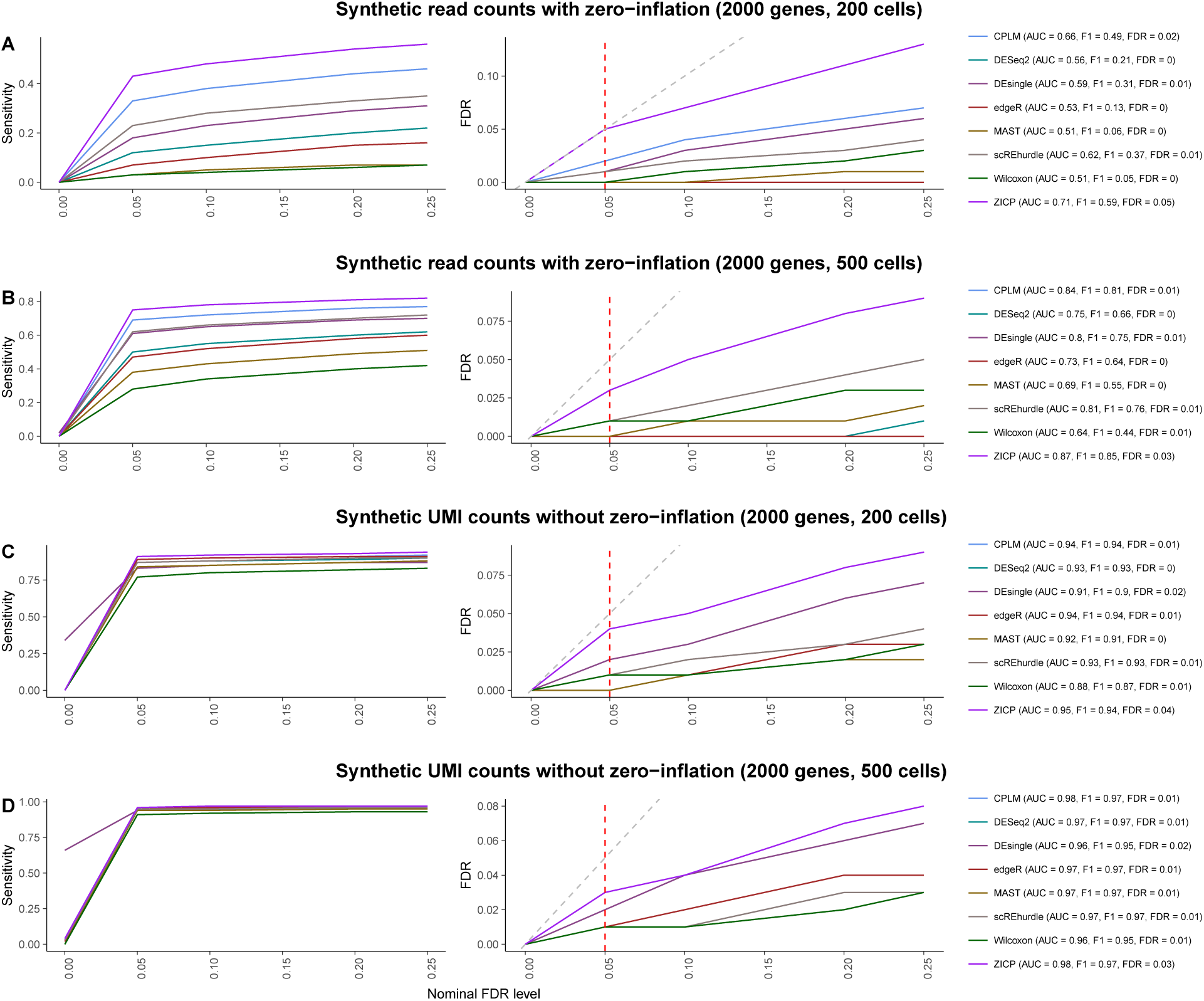
Performance results (Sensitivity and observed FDR) of Tweedieverse compared to existing DE methods using synthetic counts with and without zero-inflation for various *α*-level cutoffs. The performance of differential expression (DE) methods are compared using Splatter-simulated (**A, B**) synthetic read counts and (**C, D**) UMI counts depicting trade-offs between Sensitivity (left column) and observed FDR (right column) (*y*-axis) for various *α*-level cutoff (*x*-axis) are shown for each of the tested differential expression methods. Tweedieverse is more powerful than other methods across a range of FDR cutoffs in both read counts (**A, B**) and UMI counts (**C, D**). In the right plot, the grey line is the expected identity relationship between the observed FDR and nominal FDR level. The red line parallel to the *y*-axis is the nominal threshold for FDR in multiple testing. Values are summarized averages over 100 iterations. The area under the receiver operating characteristic (ROC) curves (AUC) and F1 scores are calculated at α = 0.05. The number of simulated genes and cells for each scenario are reported in the title of each panel. It is to be noted that due to unavoidable overlaps induced by identical FDRs at various α-levels, some of the methods including CPLM are not distinguishable in the overlaid FDR plots.

### 3.3 Re-analysis of public single-cell transcriptomics datasets

We next applied Tweedieverse to analyze two publicly available biologically heterogeneous scRNA-seq data sets. The first dataset (referred to here as the Brain data) contains expression counts of 100 single-cells generated from two different cell types: oligodentrocytes cells (*N* = 38) and astrocytes cells (*N* = 62). This non-UMI dataset is obtained from the R package *SC2P* (Wu et al., 2018), and is available from the Gene Expression Omnibus database under accession number GSE67835. We also analyzed a UMI dataset (Zheng et al., 2017) consisting of 13,713 genes and 2638 cells (referred to here as the PBMC data) collected from peripheral blood mononuclear cells (PBMCs) where we focused on comparing CD4+ T-cells (N = 271) with CD8+ T-cells (*N* = 1180) as in previous studies (Ma et al., 2020). The processed dataset is available from 10x Genomics website: https://support.10xgenomics.com/single-cell-gene-expression/datasets/1.1.0/pbmc3k. For a clear presentation of results, we consider 4 representative methods (scREhurdle, DESeq2, edgeR, MAST) along with CPLM and ZICP to examine the performance of various methods. We used the cell types provided by the authors to detect DE genes on these datasets.

In terms of the number of DE genes detected, CPLM and ZICP identified *G* = 1330 and *G* = 2057 DE genes, respectively, in the Brain dataset, while also containing a significant overlap with all other methods (*G* = 705) (**Fig. 4**). Both CPLM and ZICP uniquely detected a substantial number of DE genes (*G* = 34 and *G* = 433, respectively) that remained undetected otherwise. The same conclusion holds on the second dataset we analyzed, where ZICP detected the most unique DE genes (*G* = 168) with *G* =177 DE genes overlapping with other methods (**Fig. 5)**. Note that, not all methods successfully generated valid output in these datasets. To enable a head-to-head comparison of all six methods, we analyzed another scRNA-seq dataset of human pre-implantation embryonic cells (Petropoulos et al., 2016), where Tweedieverse remained one of the best methods along with scREhurdle with comparable number of discoveries (**Web Fig. S10**). In terms of computational time, both CPLM and ZICP scale well to large datasets although they are considerably slower than other frequentist DE approaches (**Web Table S2**). This is the price to be paid for exploring the parameter space of the index parameter p, and the computational demands of Tweedieverse is certainly less than scREhurdle, where we encountered severe convergence issues for both large and highly sparse datasets. Overall, Tweeedieverse remained stable without numerical issues across a range of sparse and non-sparse datasets.

**Figure 4.**
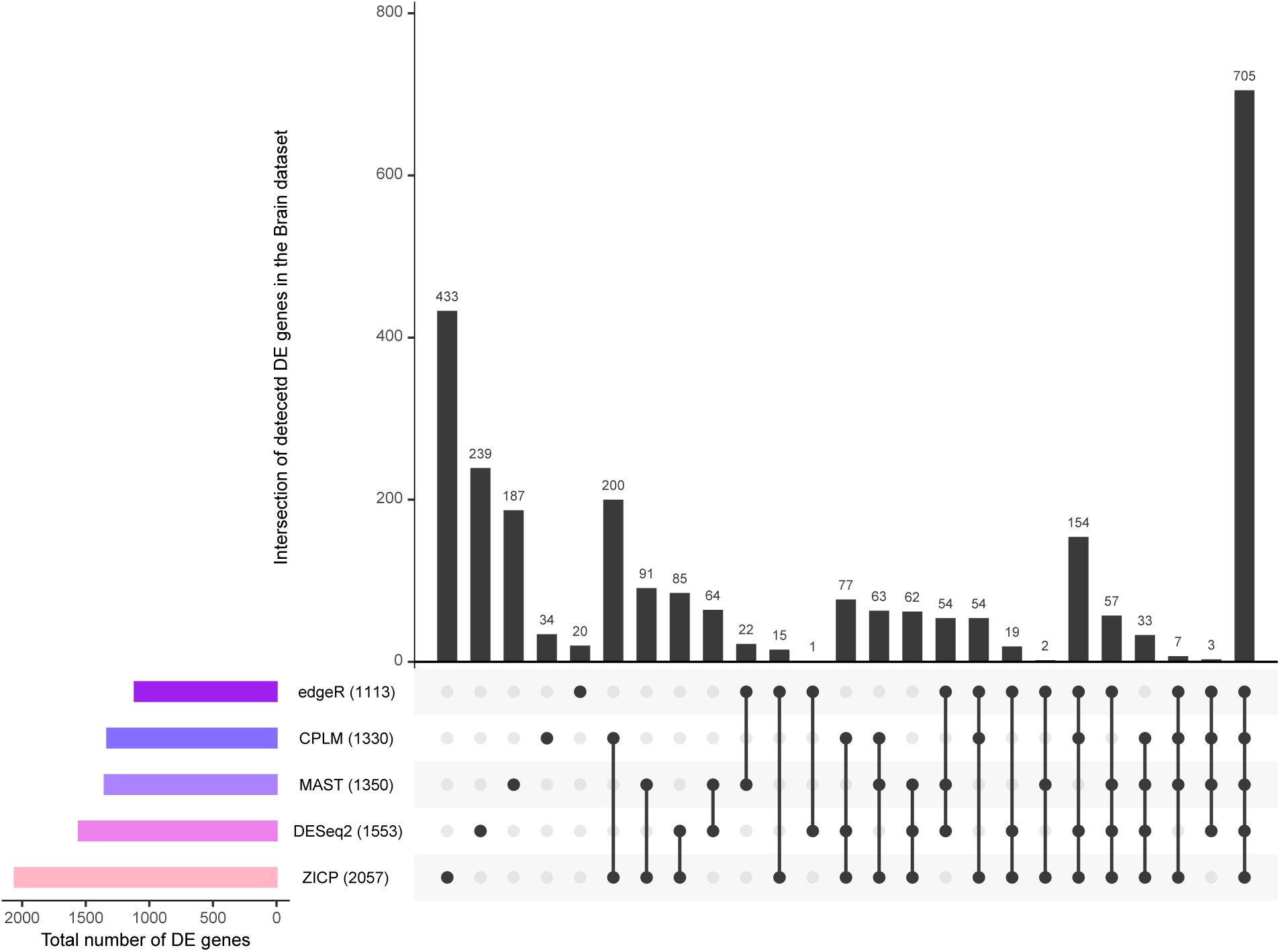
UpSet plot of number of DE genes detected across six scRNA-seq DE methods using human brain data. Using six scRNA-seq DE methods, the number of DE genes detected between astrocytes (*N* = 62 cells) and oligodendrocytes (*N* = 38 cells) groups in the human brain data (Darmanis et al., 2015). Numbers in parentheses represent the total number of DE genes identified by the corresponding method. Genes with observed FDR smaller than *α* = 0.05 are deemed as DE. We note that scREhurdle fails to execute without error for this dataset, which happens when the evidence lower bound (ELBO) stopping rule results in premature termination of the variational Bayes optimizer.

**Figure 5.**
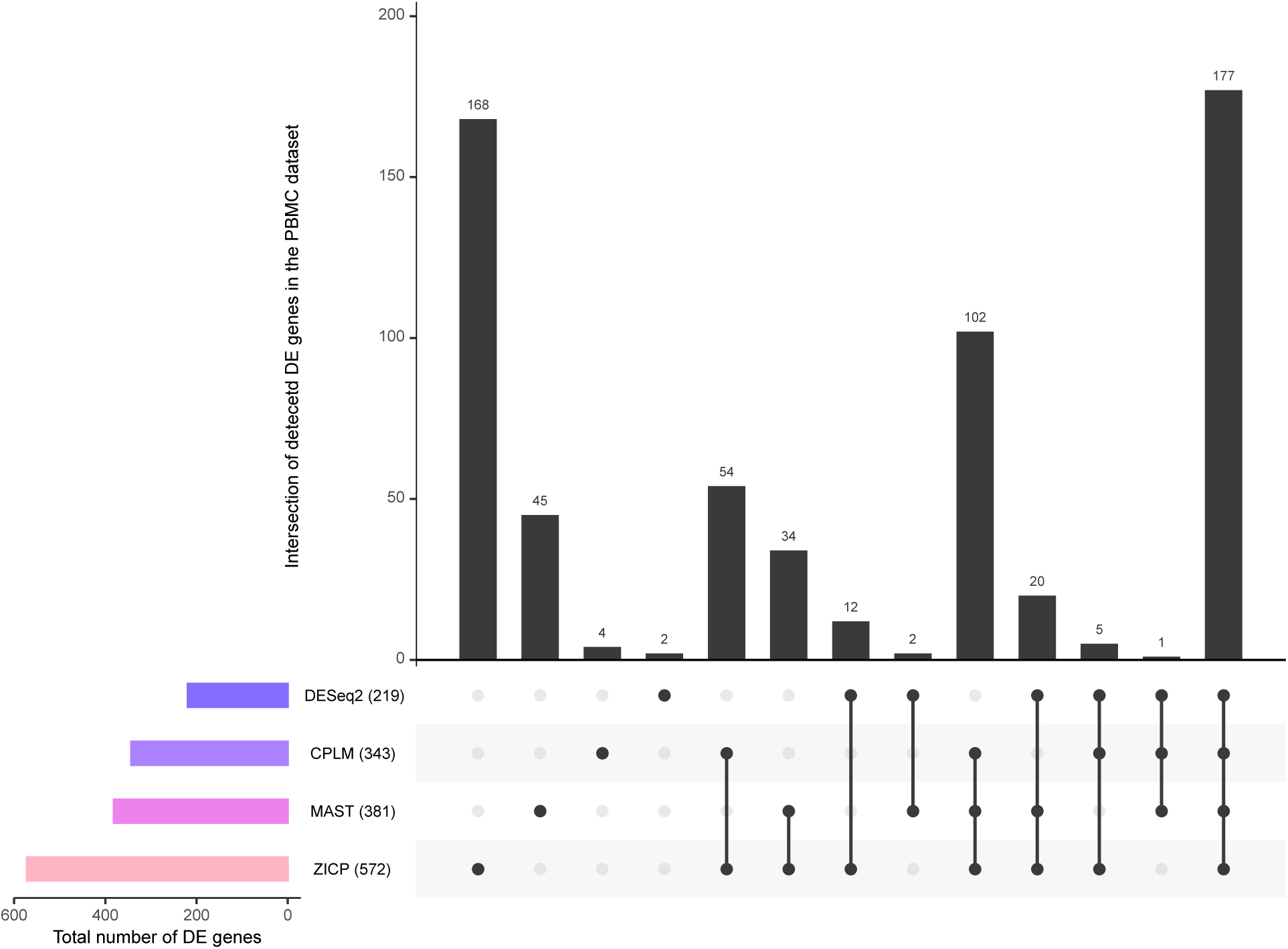
UpSet plot of number of DE genes detected across six scRNA-seq DE methods using PBMC data. Using six scRNA-seq DE methods, the number of DE genes detected between CD8+ (*N* = 271 cells) versus CD4+ T cells (*N* = 1180 cells) in the PBMC data (Zheng et al., 2017). Numbers in parentheses represent the total number of DE genes identified by the corresponding method. Genes with observed FDR smaller than *α* = 0.05 are deemed as DE. We note that scREhurdle and edgeR fail to execute without error for this dataset. For edgeR, the error happens during the *estimateTagwiseDisp* step when some of the estimated dispersions produce negative values, causing the final *glmFit* step to fail. For *scREhurdle*, the error happens when the evidence lower bound (ELBO) stopping rule results in premature termination of the variational Bayes optimizer.

One of the top hits only identified by Tweedieverse in the Brain data is the gene *AFAP1L1* (**Fig. 6A**), which has been reported to show differential expression patterns in human brain that interacts with cortactin and localizes to invadosomes (Snyder et al., 2011). In another study of spindle cell sarcoma, *AFAP1L1* has been reported as a candidate for metastasis predicting marker (Furu et al., 2011). Another gene *SERPINF1*, which is one of the top hits upregulated in oligodendrocytes has been recently shown to be one of the highly enriched genes in Oligodendrocyte progenitor cells (OPCs), which have multiple functional roles in both homeostasis and disease (Beiter et al., 2020). SERPINF1 is another name for pigment epithelium-derived factor (PEDF), which has anti-tumor effects particularly in tumors where PEDF exerts both an indirect impact on tumor angiogenesis and a direct effect on tumor cells (He et al., 2015). Such anti-tumor effects have been observed in different cancers, including gliomas. Hence, PEDF could be a potential molecular therapeutic target in anti-cancer activities (Ek et al., 2006).

**Figure 6.**
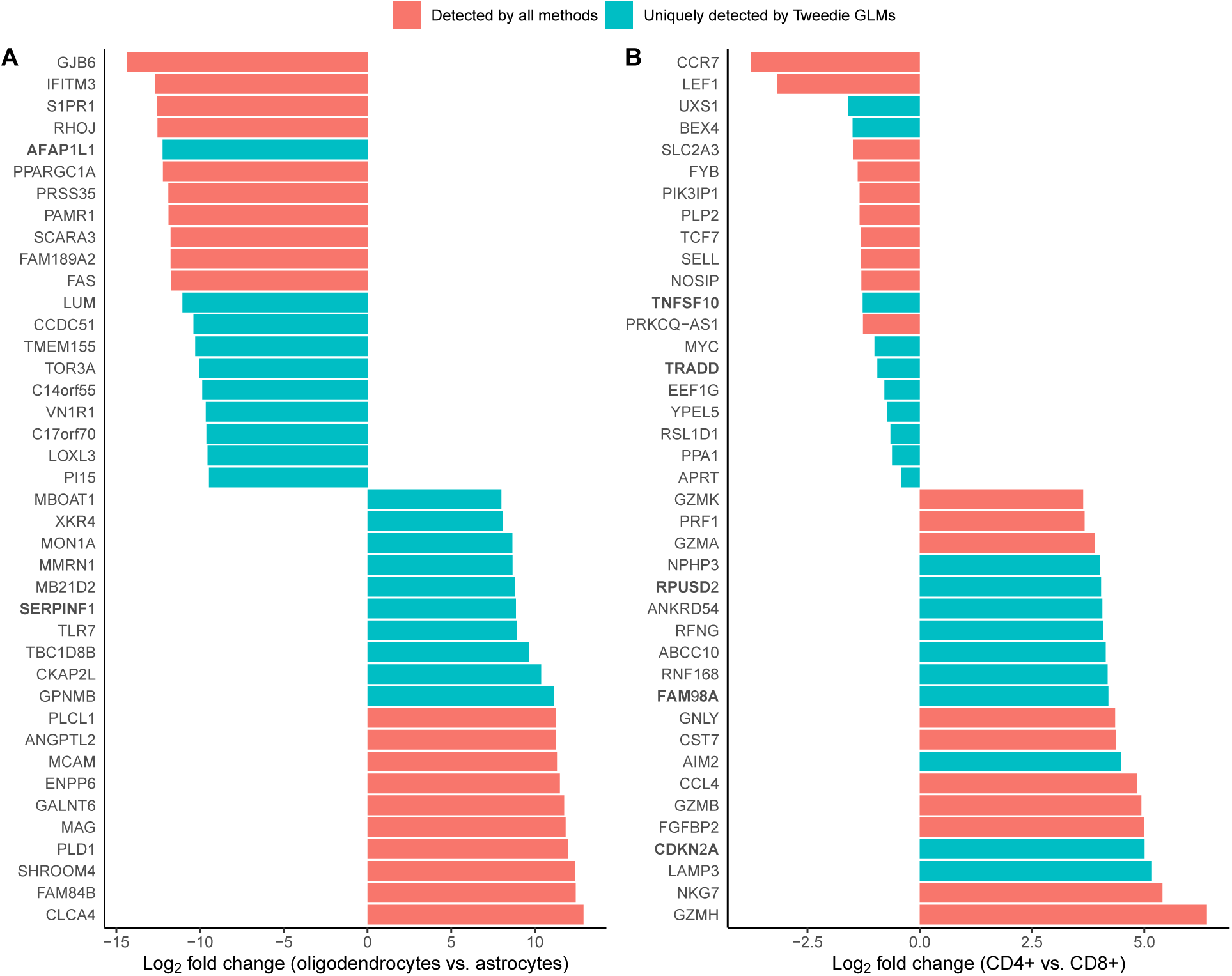
Effect sizes of DE genes uniquely identified from Tweedieverse and detected by all methods. Top 20 statistically significant DE genes (*y*-axis) (observed FDR smaller than *a* = 0.05) from (**A**) astrocytes versus oligodendrocytes comparison and (**B**) the CD4+ versus CD8+ T cells comparison. Each row illustrates a per-gene log_2_ effect size (*x*-axis). Genes are colored by two groups: (1) common genes identified by all methods in red and (2) genes only uniquely determined by the Tweedie GLMs in green (i.e. genes identified by either CPLM or ZICP or both). In addition to reporting previously reported DE genes, Tweedieverse is able to detect genes with biologically meaningful small effect sizes not detected by existing methods. Specific biologically relevant candidate genes deserving follow-up experiments (as described in Section 3.3) are highlighted in bold.

Among the top genes detected in the PBMC dataset, most of the DE genes have small effect sizes, which is consistent with the notion that the two examined cell types (CD4+ T-cells and CD8+ T-cells) are strikingly similar, making the DE analysis difficult with many small effects (Ma et al., 2020). Following Ma et al. (2020), we also conducted a gene set enrichment analysis (GSEA) on the small set of curated 144 gene sets that contain important CD4+ and CD8+ cell type signatures (Aran et al., 2017). GSEA analysis on the unique genes detected by the Tweedie models revealed that most of the top enriched gene sets are relevant to CD4 or CD8 cell functions (**Web Fig. S11**) and several of these genes (*TNFSF10*, *CDKN2A*, *RPUSD2*, *FAM98A*, *TRADD*) are associated with multiple immunologic signatures (**Fig. 6B**). Interestingly, a similar GSEA analysis on the common genes found by all methods did not yield any significant results. These findings thus suggest that Tweedieverse not only identifies more DE genes and more enriched gene sets, but also most of these uniquely detected genes have the potential to reveal important functional mechanisms, which can be missed by existing methods.

## 4 Discussion

We have described a series of statistical methods to detect DE features in discrete scRNA-seq data based on a three-parameter Tweedie family of distributions. Our per-gene estimation approach affords the Tweedie model a great deal of flexibility and adaptivity to different sparsity and zero-inflation levels and experimental platforms, ensuring robustness and applicability across a wide range of scRNA-seq expression profiles. Although univariable differential expression analysis between homogeneous cell populations is the use case of our method, mixed effects and multivariable extension of our approach to incorporate cell-to-cell heterogeneity and multiple covariates is straightforward. Additionally, beyond scRNA-seq studies, our method is extensible to alternative data modalities such as metagenomics and metatranscriptomics with similar zero-inflated and semi-continuous count data distributions (Mallick et al., 2017). We conjecture that due to their flexibility, our Tweedie models would retain their strong empirical and theoretical properties in these other settings.

While our proposed methodology is comparable in spirit to that of existing approaches, our method addresses several methodological challenges not addressed by previous scRNA-seq DE methods. For example, despite a large number of developments in the area, a lack of interoperability between platform-specific methods has been a major concern and no comparable framework offers the flexibility of fitting a variety of regression models (appropriate for both UMI counts and read counts) under one estimation umbrella. Owing to the flexibility of the Tweedie formulation, we address this by providing users a variety of options including count, continuous, and zero-modified regression models based on the CPLM and ZICP models.

From a practical standpoint, our method offers a comprehensive and general solution for gene-level analysis of scRNA-seq data. While the standard Tweedie model (CPLM) is well-suited for sparse UMI counts, the zero-inflated counterpart of the Tweedie model (ZICP) represents an attractive alternative for zero-inflated read counts. With a computationally efficient algorithm based on the profile likelihood and automated index parameter estimation, our method is also scalable to datasets with tens of thousands of genes measured on tens of thousands of samples. The self-adaptive nature of the Tweedie model enables an automatic tuning of the index parameter (*p*), thus avoiding the need to specify a grid of plausible values by the user. Finally, the encapsulation of these strategies in the paradigm of GLMs enables the treatment of both simple and complex experimental designs, making it applicable to a variety of general epidemiological studies and controlled trials.

In addition to the versatility of the proposed methods, our extensive simulation studies and real data analyses reveal that both the CPLM and ZICP methods perform as well as or better than existing methods in controlling the FDR, while consistently maintaining superior power across diverse scRNA-seq datasets (both simulated and real). Future work could provide several refinements to the method. Perhaps most importantly, it may be possible to borrow information across genes to obtain shrinkage estimates that can facilitate a better ranking of the DE genes based on regularized log fold changes (Stephens, 2017; Zhu et al., 2019). In a similar vein, the CPLM model can be seamlessly integrated into the hurdle model framework to improve upon existing methods (i.e. MAST and scREhurdle) without additional computational overhead. Finally, the ZICP model can be marginalized to provide a direct interpretation of the overall effect of the covariates on the marginal mean (Preisser et al., 2016; Long et al., 2014). We thus aim for our method to provide a happy medium, capable of serving as a one-stop-shop for DE analyses for a wide range of scRNA-seq data types.

## Acknowledgements

SC was supported by the Intramural Research Program of *Eunice Kennedy Shriver* National Institute of Child Health and Human Development (NICHD) of the National Institutes of Health (NIH). AR was supported by the National Science Foundation (NSF) grant DEB-2028280 and the Bill and Melinda Gates Foundation grant INV-016930. SCH was supported by the National Human Genome Research Institute (NHGRI) of the National Institutes of Health (NIH) under the award number R00HG009007.

## Implementation and Software Availability

The implementation of the associated software (*Tweedieverse*) is publicly available with source code, documentation, tutorial, and as an R package at https://github.com/himelmallick/Tweedieverse. Analysis scripts for the synthetic benchmarking (as described in Section 3.2) are available at https://github.com/himelmallick/BenchmarkSingleCell.

