## Supporting Information for "Differential expression of single-cell RNA-seq data using Tweedie models"

Mallick et al.

#### 1 Supporting Text

##### Web Appendix S1: Connection with exponential dispersion models

The Tweedie distribution belongs to a subclass of the exponential dispersion model (EDM) family defined by

$$f(y|\theta,\phi) = a(y,\phi)\exp\left[\frac{y\theta - \kappa(\theta)}{\phi}\right],$$

where $a(\cdot)$ and $\kappa(\cdot)$ are known normalizing functions, $\theta$ is the natural parameter, and $\phi > 0$ is the dispersion parameter (Zhang, 2013). The mean $\mu$ and variance $\sigma^2$ of EDM are given by $\mu = \kappa'(\theta)$ and $\sigma^2 = \phi\kappa''(\theta)$. Due to the dependence between $\theta$ and $\mu$, $\kappa''(\theta)$ can be characterized by the variance function, $\kappa''(\theta) = V(\mu)$, leading to a direct mean-variance relationship given by $\sigma^2 = \phi V(\mu)$. The Tweedie family of distributions corresponds to a special power variance function given by $V(\mu) = \mu^p$, $p$ being the power parameter.

When $1 < p < 2$, the Tweedie distribution is characterized by a mixture of Poisson and a compound sum of i.i.d. Gamma random variables, i.e., $Y = \sum_{i=1}^{T} X_i$ and $T \sim Pois(\lambda)$, where $X_i \overset{iid}{\sim} G(\alpha_1, \alpha_2)$ and $T \perp X_i$, $\forall i$. Therefore, the joint distribution of $Y > 0$ and $T$ can be expressed as

$$f(y,t|\lambda,\alpha,\beta) = \frac{\lambda^t \exp(-\lambda)}{t!}\frac{y^{t\alpha_1 - 1}\exp(-y/\alpha_2)}{\alpha_2^{t\alpha_1}\Gamma(t\alpha_1)}, \tag{1}$$

where the marginal distribution of $Y$ can be obtained by integrating out $T$ from (1). If $T = 0$, then $Y = 0$ and the Tweedie distribution has probability mass of the form $\exp(-\lambda)$ at the origin. If $T > 0$, $Y$ can be characterized by the sum of $T$ i.i.d. Gamma random variables. Following Kurz (2017), the parameters of the Tweedie distribution are directly related to the EDM parameters as follows: $\lambda = \frac{\mu^{2-p}}{\phi(2-p)}$, $\alpha_1 = \frac{2-p}{p-1}$, and $\alpha_2 = \phi(p-1)\mu^{p-1}$. The normalizing constant $a(y, \phi)$ does not have a closed form but can be approximated by Stirling's formula for the Gamma function and a Fourier inversion method for the infinite series (Kurz, 2017).

#### Web Appendix S2: Maximum likelihood estimation

For carrying out statistical inference, we denote the $n$ observations (cells) of the count response (i.e. per-gene expression) as $\boldsymbol{Y} = (Y_1, \ldots, Y_n)$, where $Y_i \sim Tw\,(\mu, \phi, p)$ for UMI counts, or $Y_i \sim ZITw\,(\mu, \phi, p)$ for read counts. For a known value of the index parameter $p$, the Tweedie model, due to its connection with the exponential dispersion family as described before, can be estimated using existing inferential procedures such as Fisher's scoring algorithm (McCullagh and Nelder, 1989). However, in most situations, when the parameter $p$ is not known beforehand, a profile likelihood approach (Cox and Reid, 2017) can be considered to estimate the parameters. It is to be noted that, due to the dependency of the normalizing constant $a\,(y, \phi, p)$ on the index parameter $p$, the corresponding objective function is intractable as the Tweedie density function has no closed form expression for $1 < p < 2$ (Zhang, 2013). Thus, numerical methods such as Fourier inversion strategy (Dunn and Smyth, 2007) must be employed for evaluating the Tweedie density function. Once evaluated, an estimate of $\theta$ can be obtained by numerically optimizing the log of the profile likelihood $l\left(\theta|\boldsymbol{y}, \hat{\boldsymbol{\beta}}(\theta)\right)$ subject to the constraints $\phi > 0$ and $p \in (1, 2)$. Specifically, $\theta = (\phi, p)$ can be estimated by profiling out $\beta$ and maximizing $l(\theta)$ as

$$\hat{\theta} = \underset{\theta}{\operatorname{argmax}}\; l\left(\theta|\boldsymbol{y}, \hat{\boldsymbol{\beta}}(\theta)\right), \tag{2}$$

where for a given estimate of $\theta$, the regression coefficients $\boldsymbol{\beta}$ are estimated as $\hat{\boldsymbol{\beta}}(\hat{\theta})$ using the scoring algorithm mentioned before. This process of estimating $\theta$ by profiling $\boldsymbol{\beta}$ out of the log-likelihood and updating $\hat{\boldsymbol{\beta}}$ based on the current $\hat{\theta}$ is continued until convergence.

#### Web Appendix S3: The EM algorithm

For the zero-inflated Tweedie model, we employ an Expectation-Maximization (EM) algorithm (Dempster et al., 1977) detailed below subject to the constraint $\phi > 0$ and $p \in (1, 2)$ and maximize the following objective function

$$l(\Theta|\boldsymbol{y}) = \sum_{i=1}^{n} \log[q_i * 1(y_i = 0) + (1 - q_i) * Tw(y_i, \beta, \phi, p)]. \tag{3}$$

We formulate Equation (3) as a 'missing data' problem and introduce a set of latent variables $z_i$, where $z_i = 1$ if $y_i$ is generated from the 'zero' component and $z_i = 0$ if $y_i$ comes from the Tweedie component. The 'complete data' can be expressed with $(y_i, z_i)$, $i = 1, \ldots, n$, where we only observe $y_i$'s and treat $z_i$'s as 'missing data', leading to the following log-likelihood:

$$l(\Theta|\boldsymbol{y}, \boldsymbol{z}) = \sum_{i=1}^{n} [z_i Z_i \gamma - \log(1 + \exp(Z_i \gamma) + (1 - z_i) \log(Tw(y_i, \beta, \phi, p))]. \tag{4}$$

Starting with an initial guess of the coefficient estimates, the algorithm iterates between $E$ step and $M$ step, where at the $E$ step of the algorithm, we calculate the expectation of the log-likelihood of the 'complete data' by replacing $z_i$'s by their conditional expectations given the current estimates. At the $M$ step, we estimate the coefficients by maximizing the 'complete data' log-likelihood in Equation (4) using the profile likelihood approach. To summarize, the algorithm proceeds as follows:

1. Start with an initial guess of the estimate of the coefficients.

2. $E$ step: At the $t^{\text{th}}$ iteration, calculate

$$z_i^{(t)} = \begin{cases} \left[1 + \exp\left(-(\lambda + \boldsymbol{X}_i\beta)\right)\right]^{-1}, & \text{if } y_i = 0, \\ 0, & \text{if } y_i > 0, \end{cases}$$

   where $\lambda = \frac{\exp((2-p)\boldsymbol{X_i}\beta)}{(2-p)\phi}$.

3. $M$ step: Maximize the objective function $Q(\beta, \gamma|\beta^t, \gamma^t)$ which can be expressed as a sum of logistic log-likelihood and a weighted Tweedie log-likelihood and thus can be separately optimized to estimate $\boldsymbol{\beta}$ and $\boldsymbol{\gamma}$, where $Q(\beta, \gamma|\beta^t, \gamma^t) = \sum_{i=1}^{n}[z_i^{(t)} Z_i\gamma - \log(1+\exp(Z_i\gamma)] + (1 - z_i^{(t)}) \log(g(y_i : \beta, \phi, p))$. We use the profile likelihood approach described before to optimize the second component and the functionalities of the logistic GLM to optimize the first component.

4. Repeat 2 and 3 until convergence, $t = 1, 2, \ldots$

### 2 Supporting Tables

**Web Table S1**: **List of datasets used in Figure 1 and the rest of the manuscript**. Datasets are broadly categorized by (i) measurement type (UMI counts or read counts) and (ii) experiment type (negative control and real-world). All the negative control datasets are available from Svensson (2020). The Islam et al. (2011) dataset is available from the Gene Expression Omnibus database under accession number GSE29087, whereas the Brain dataset (Darmanis et al., 2015) is obtained from the Gene Expression Omnibus database under accession number GSE67835. The PBMC dataset is available at 10x Genomics website (`https://support.10xgenomics.com/single-cell-gene-expression/datasets/1.1.0/pbmc3k`). Rest of the datasets are available from `https://github.com/gongx030/scDatasets`. Sparsity refers to the percentage of zero counts in the expression matrix; in particular, for an expression matrix $X$ with $N$ cells and $M$ genes, we calculate sparsity as $100 * (N * M - \text{sum}(X > 0))/(N * M))$.

| Dataset | Count | Experiment | No. of cells | No. of genes | Sparsity |
|---|---|---|---|---|---|
| Svensson et al. (2017) Dataset 1 | UMI | Negative control | 2000 | 24116 | 97.4 |
| Svensson et al. (2017) Dataset 2 | UMI | Negative control | 2000 | 24116 | 97.6 |
| Zheng et al. (2017) Dataset 1 | UMI | Negative control | 1015 | 92 | 47.2 |
| Macosko et al. (2015) | UMI | Negative control | 84 | 959 | 73.7 |
| Klein et al. (2015) | UMI | Negative control | 953 | 25435 | 61.8 |
| Zheng et al. (2017) Dataset 2 (PBMC) | UMI | Real-world | 1180 | 13097 | 93.8 |
| Blakeley et al. (2015) | Read | Real-world | 30 | 11900 | 46.0 |
| Deng et al. (2014) | Read | Real-world | 286 | 12933 | 37.4 |
| Guo et al. (2015) | Read | Real-world | 315 | 12246 | 46.6 |
| Gong et al. (2018) | Read | Real-world | 290 | 9075 | 52.0 |
| Loh et al. (2016) | Read | Real-world | 429 | 12234 | 31.6 |
| Petropoulos et al. (2016) | Read | Real-world | 1529 | 13930 | 32.8 |
| Pollen et al. (2014) | Read | Real-world | 272 | 10688 | 50.4 |
| Islam et al. (2011) | Read | Real-world | 92 | 11796 | 64.8 |
| Darmanis et al. (2015) (Brain) | Read | Real-world | 100 | 10483 | 64.9 |

**Web Table S2**: **Running times (in minutes) of each method for each case study**. Methods are sorted approximately in increasing order of computing time. The number of cells, genes, and sparsity levels of these datasets are reported in **Web Table S1**. Methods that failed generate a valid output are reported with a dash. For edgeR, the error happens during the *estimateTagwiseDisp* step when some of the estimated dispersions produce negative values, causing the final *glmFit* step to fail. For *scREhurdle*, the error happens when the evidence lower bound (ELBO) stopping rule results in premature termination of the variational Bayes optimizer. Methods were run in R (version 3.6.3) independently using a single core of a system with an Intel Core i9 processor (2.3 GHz) and 16 GB of RAM.

| | Brain | PBMC | Petropoulos et al. (2016) |
|---:|---|---|---|
| edgeR | 0.36 | - | 0.86 |
| MAST | 1.17 | 3.37 | 1.88 |
| DESeq2 | 1.53 | 4.26 | 1.09 |
| CPLM | 14.33 | 48.81 | 15.92 |
| ZICP | 33.95 | 34.52 | 31.9 |
| scREhurdle | - | - | 91.83 |

#### 3 Supporting Figures

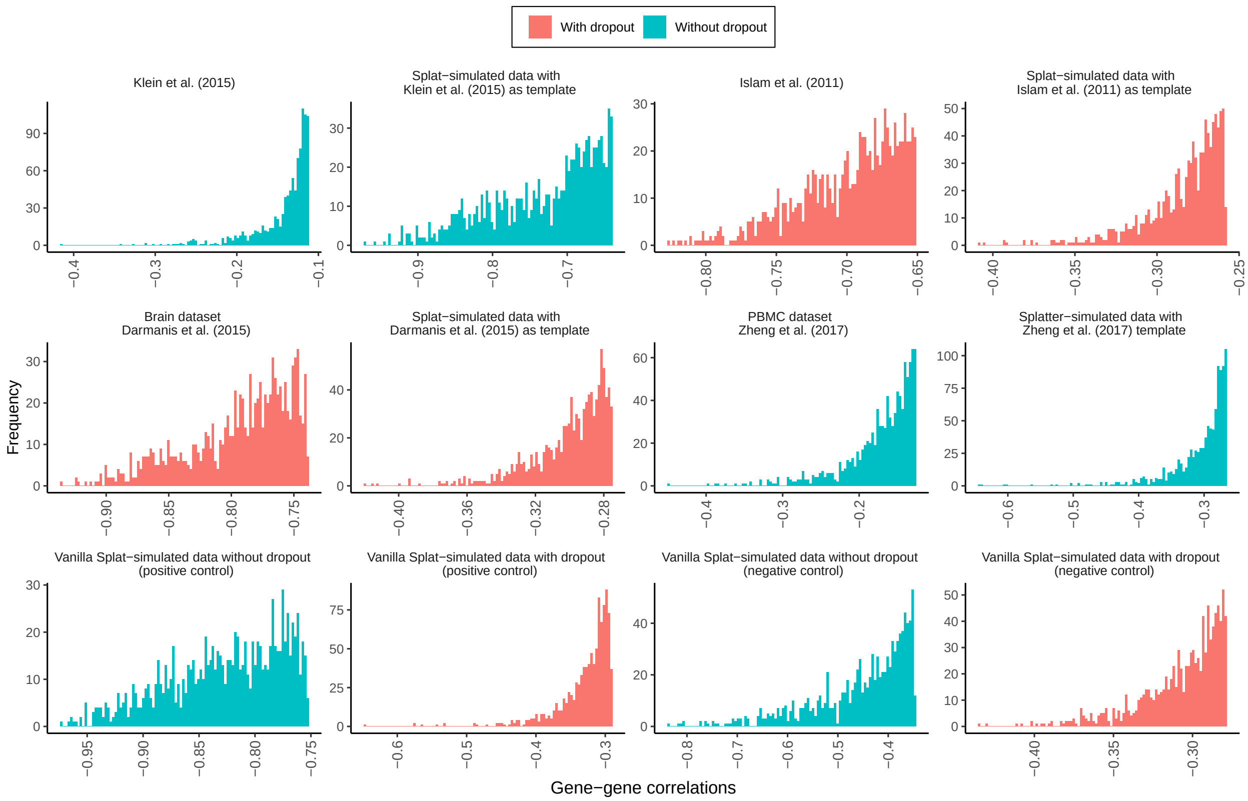


**Web Figure S1**: **Gene-gene Pearson correlations in real and simulated scRNA-seq datasets**. We used the Splatter framework (Zappia et al., 2017) for simulating synthetic scRNA-seq counts using a subset of template datasets described in **Web Table S1**. In each instance of the Splatter-simulated datasets, 2000 genes and 100 cells are generated. The effect size parameter (*de.facLoc*) is set to 1 for all the simulated datasets except for the negative control ones where it is set to zero. For visualization purposes, only top 1000 most variable genes are plotted.

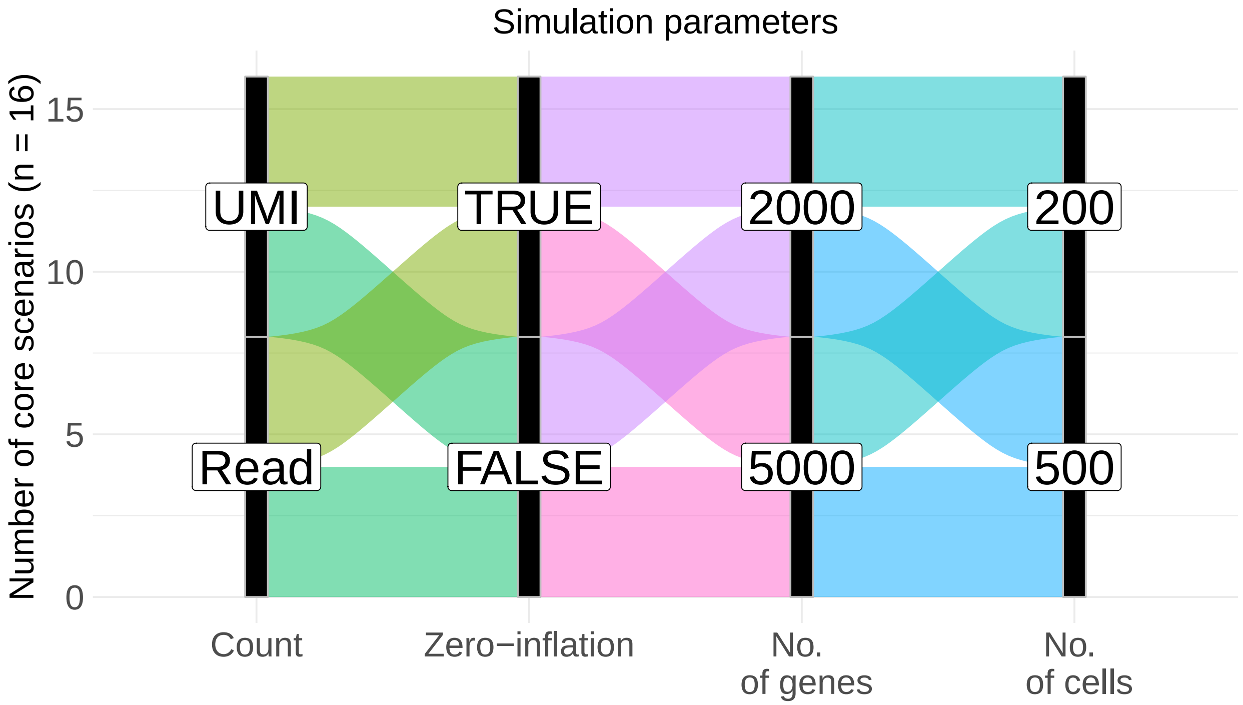


**Web Figure S2**: **Details of simulation parameters**. Two broad measurement types commonly encountered in scRNA-seq studies (UMI counts and read counts) with and without zero-inflation for varying number of cells and genes are considered leading up to 16 core scenarios. For each simulation scenario, a total of 100 datasets are generated, and within each dataset, the effect sizes ($\log_2$-fold changes) are varied between - 3 and 3 (as shown in **Web Figure S3**). The probability of a gene being DE was set to 0.1, and cells had equal probabilities of being assigned to one of the two groups. We use the Islam et al. (2011) dataset as a template for the read count simulation, whereas, the Klein et al. (2015) dataset is used as a template for generating the UMI counts.

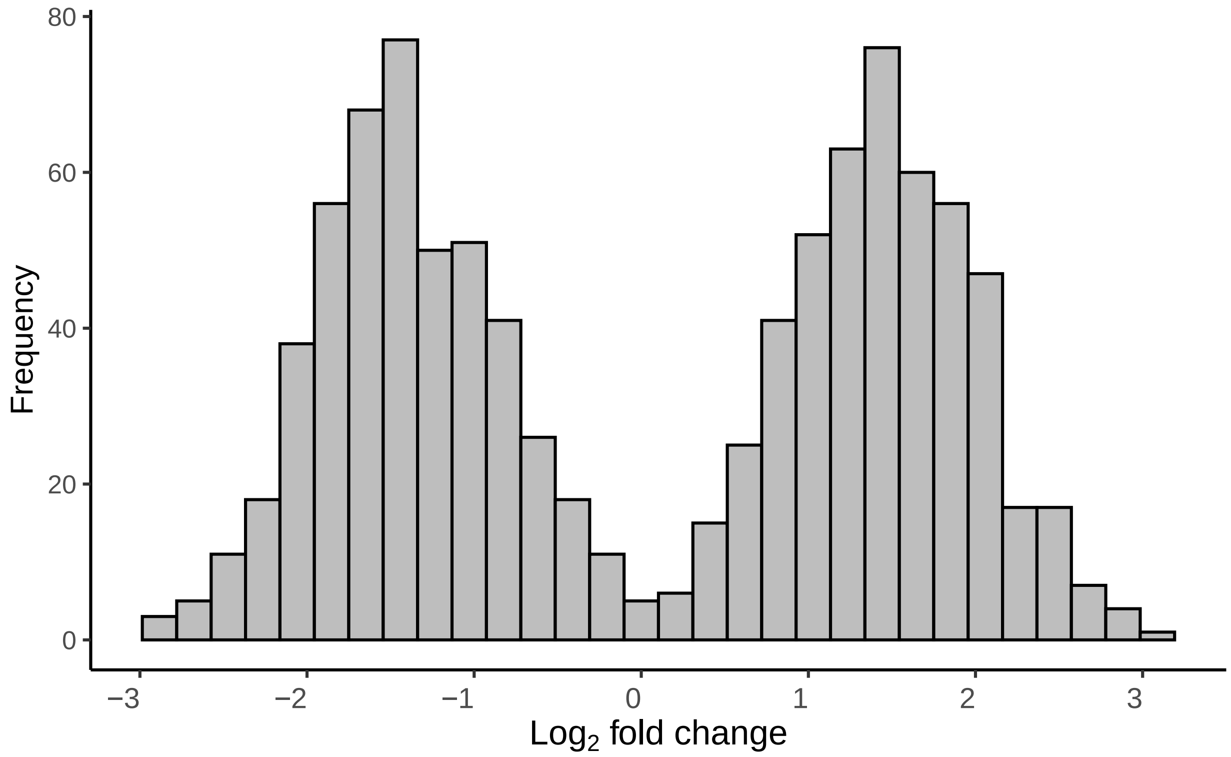


**Web Figure S3**: **The distribution of the** $\log_2$**-fold changes among DE genes in a representative Splatter-simulated dataset.** The output of Splatter contains differential expression factors (*DEFac[Group]*) showing whether a gene has differential expression (factor different from 1) or not (factor = 1) in each group. To settle on an appropriate value of the *de.facLoc* parameter, we considered several candidate values but ultimately settled on *de.facLoc* = 1. To justify our choice, we calculated the $\log_2$-fold changes for each DE gene as the $\log_2$-transformed ratio of these factors, which revealed that Splatter-generated effect sizes ($\log_2$-fold changes) realistically vary between - 3 and 3 for this choice, representing both modest (e.g. $<$ 2-fold differences) and strong (e.g. 8-fold) effect sizes.

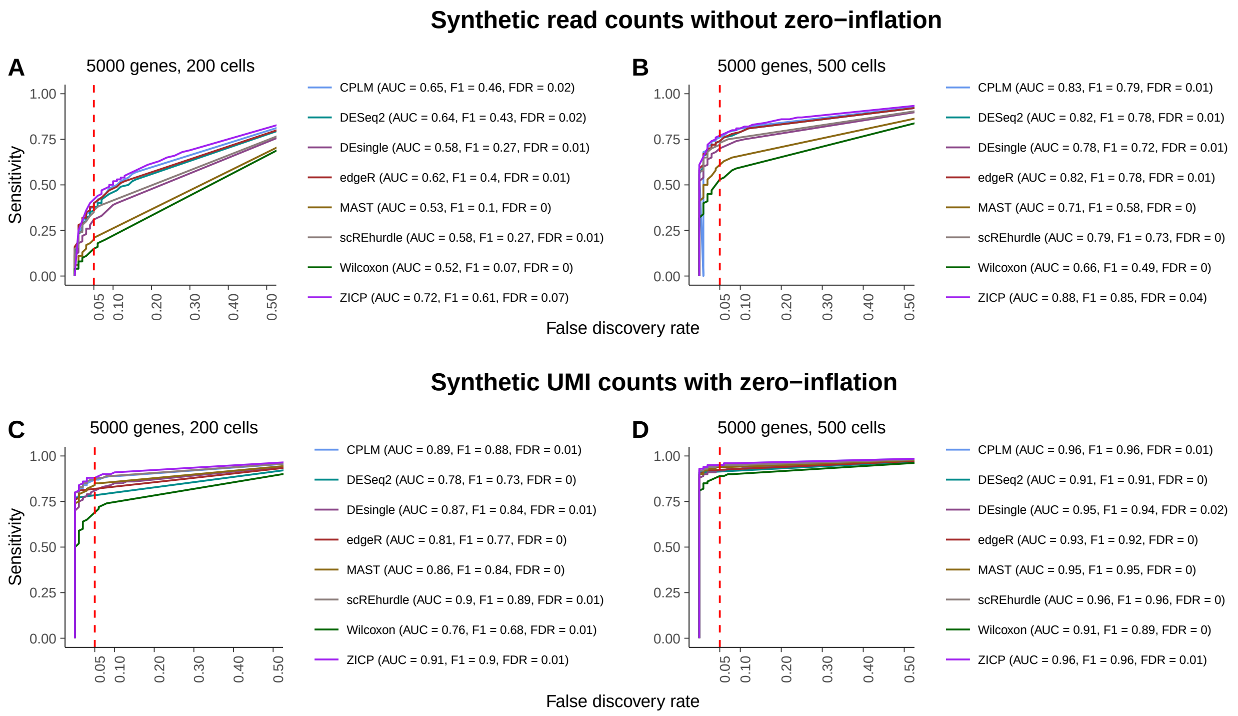


**Web Figure S4**: Similar to Figure 2 which used $G = 2000$ genes in the main manuscript, but here for datasets with larger number of genes ($G = 5000$).

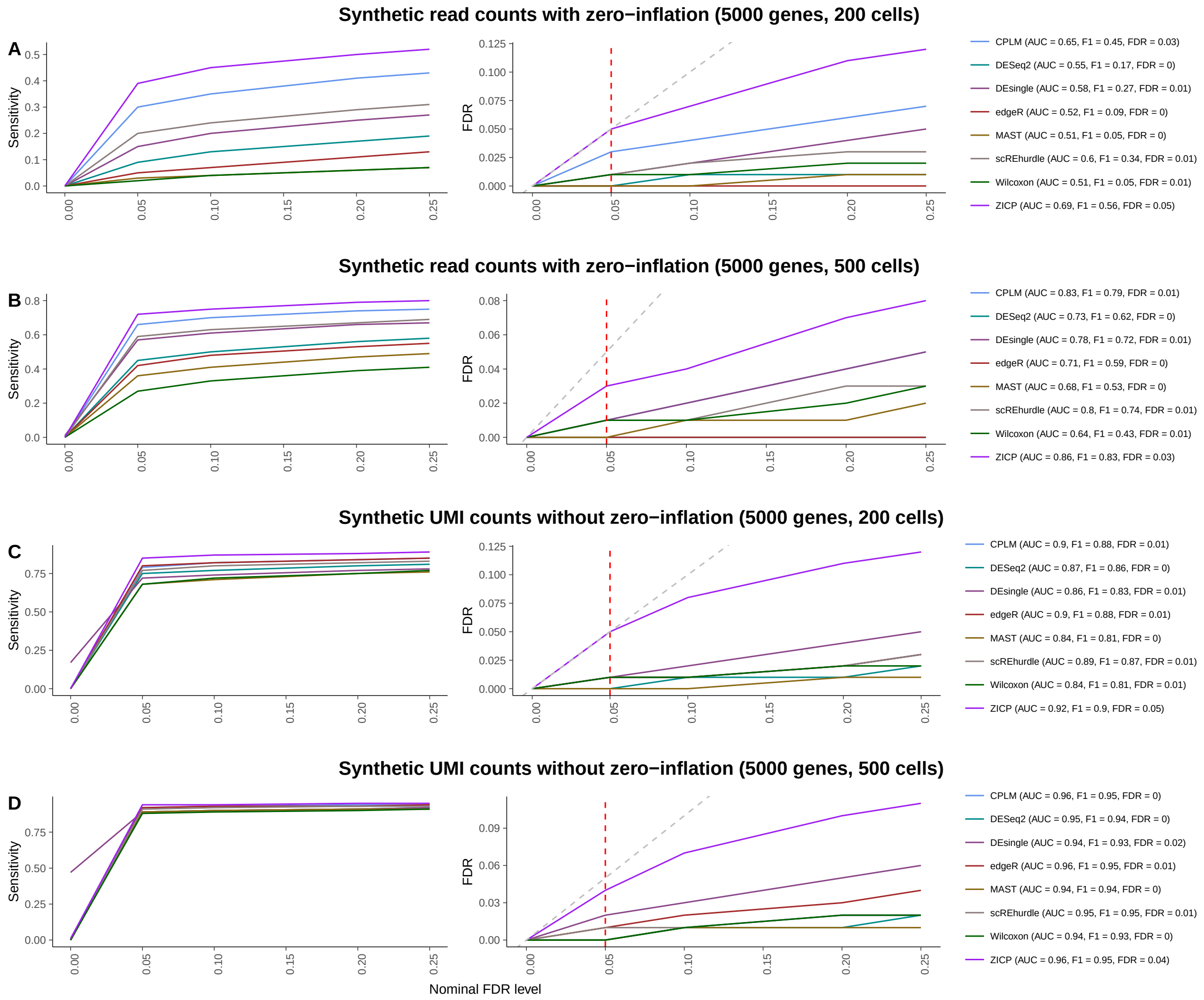


**Web Figure S5**: Similar to Figure 3 which used $G = 2000$ genes in the main manuscript, but here for datasets with larger number of genes ($G = 5000$).

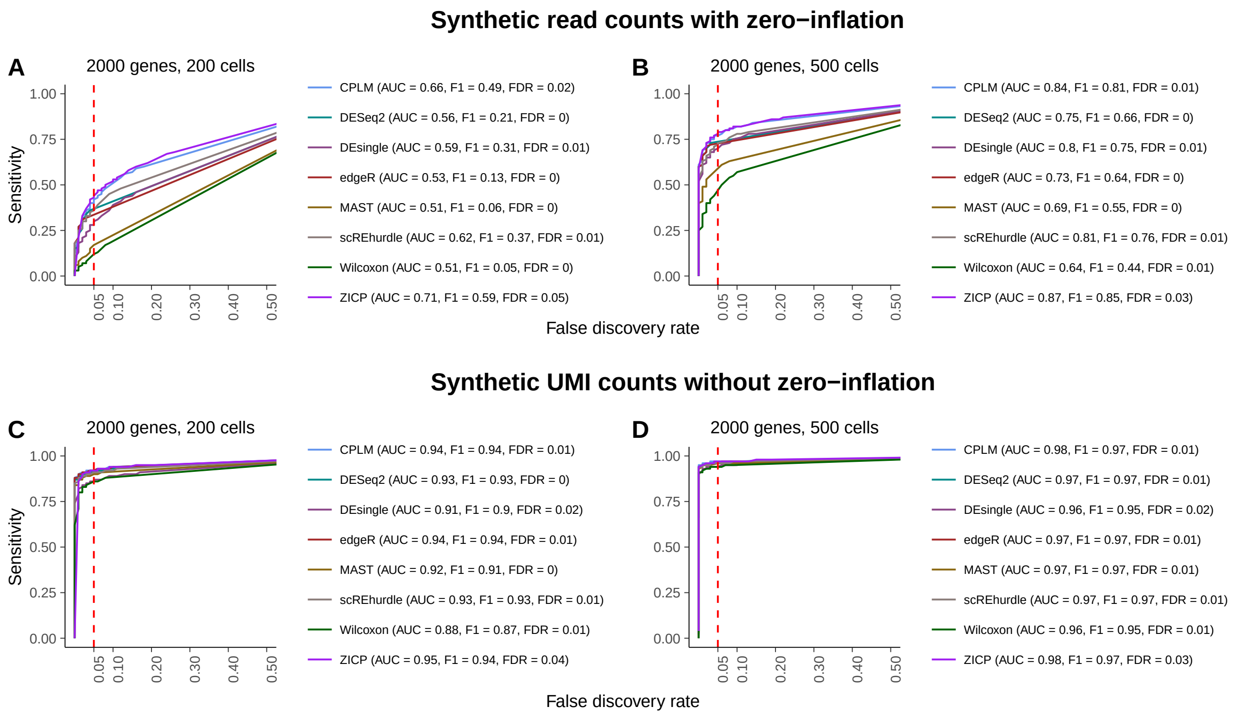


**Web Figure S6**: Similar to Figure 2 which used synthetic read counts with zero-inflation and synthetic UMI counts without zero-inflation in the main manuscript, but here considering alternative zero-inflation modes. Specifically, we consider synthetic read counts without zero-inflation and synthetic UMI counts with zero-inflation.

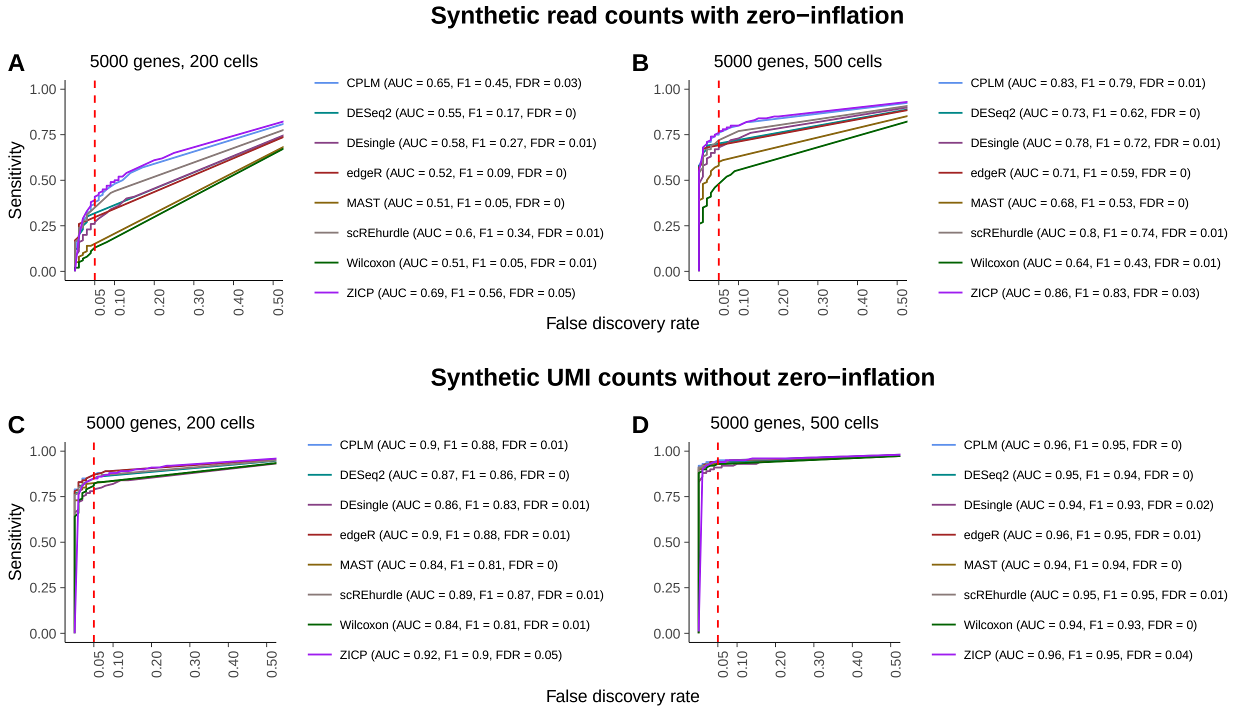


**Web Figure S7**: Similar to Web Figure S6 with synthetic read counts without zero-inflation and synthetic UMI counts with zero-inflation (which used $G = 2000$ genes), but here for datasets with larger number of genes ($G = 5000$).

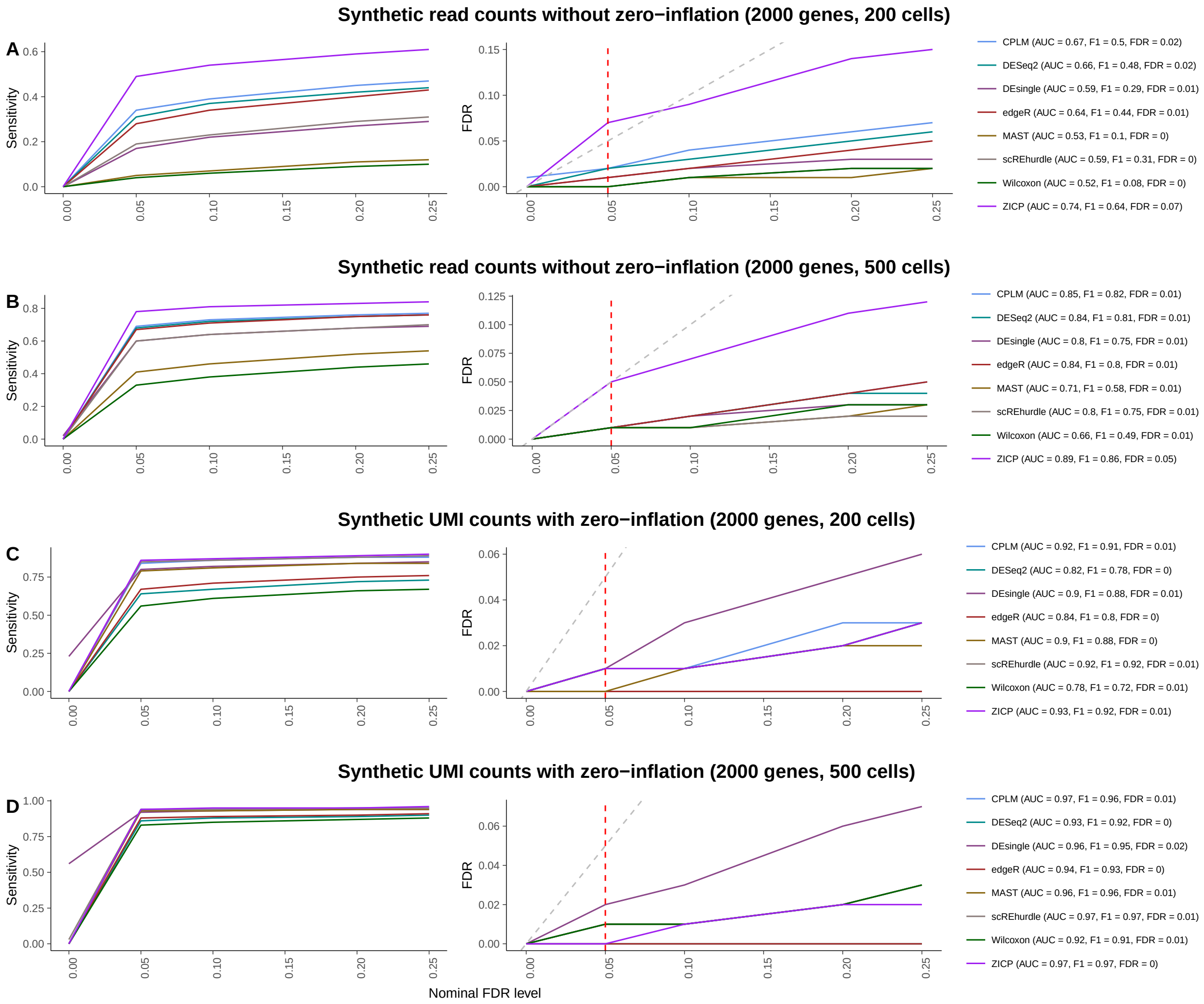


**Web Figure S8**: Similar to Figure 3 which used synthetic read counts with zero-inflation and synthetic UMI counts without zero-inflation in the main manuscript, but here considering alternative zero-inflation modes. Specifically, we consider synthetic read counts without zero-inflation and synthetic UMI counts with zero-inflation.

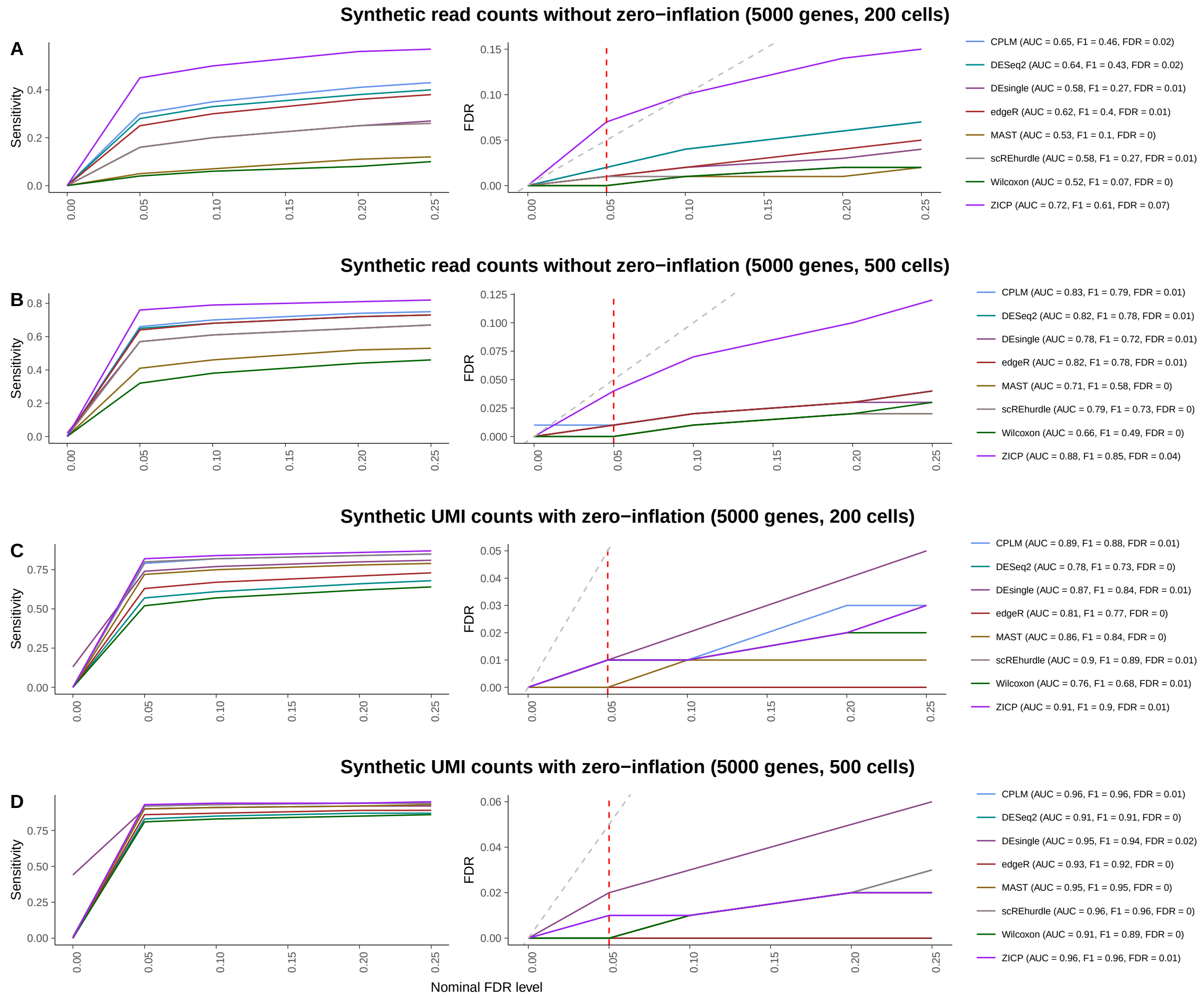


**Web Figure S9**: Similar to Web Figure S8 with synthetic read counts without zero-inflation and synthetic UMI counts with zero-inflation (which used $G = 2000$ genes), but here for datasets with larger number of genes ($G = 5000$).

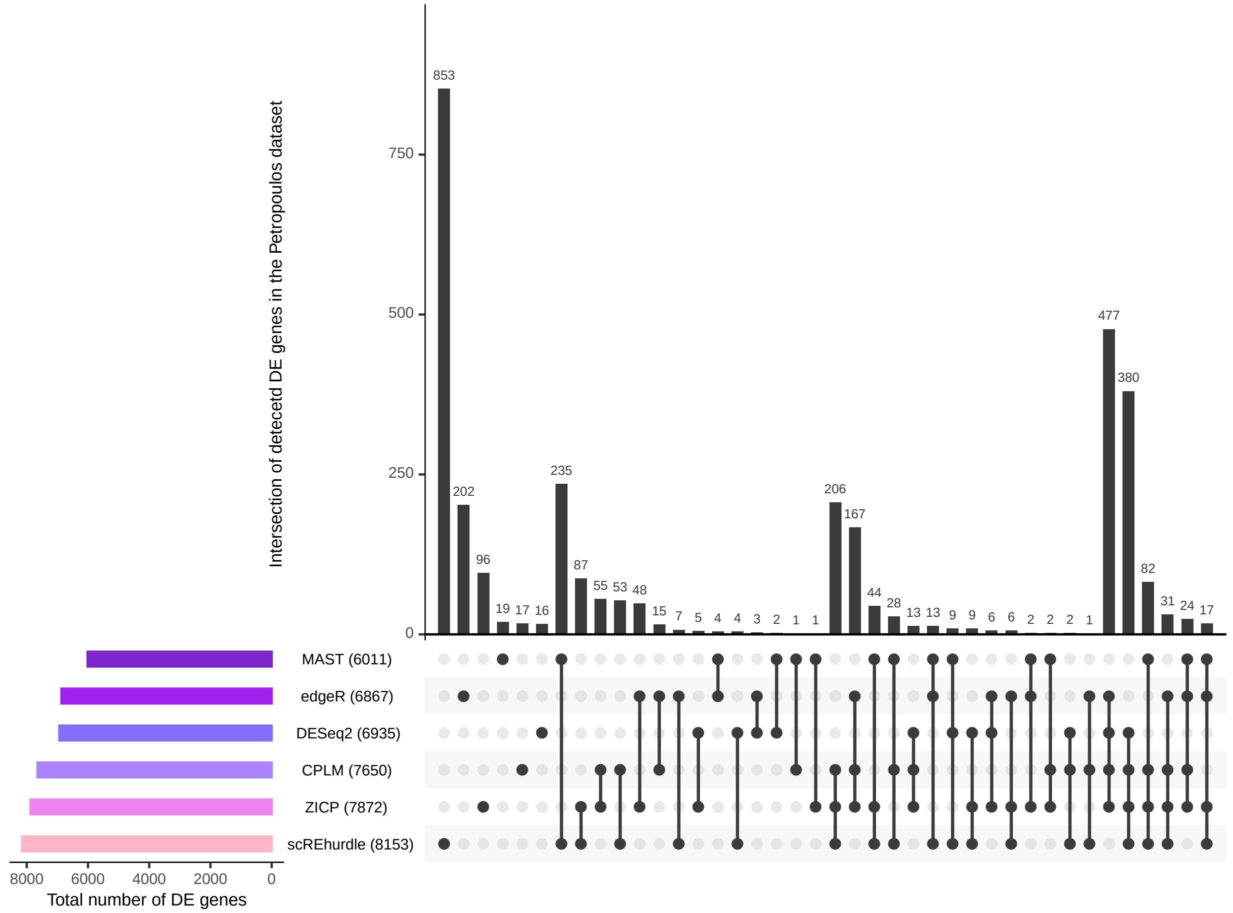


**Web Figure S10**: **UpSet plot of number of DE genes detected across six scRNA-seq DE methods using Petropoulos et al. (2016) dataset.** Using six scRNA-seq DE methods, the number of DE genes detected between embryonic day 3 (E3) ($N = 81$ cells) versus embryonic day 4 (E4) ($N = 190$ cells) groups. This dataset has recently been analyzed by Miao et al. (2018). We included this additional dataset to showcase a head-to-head comparison with scREHurdle which crashed in a majority of datasets we tested. Numbers in parentheses represent the total number of DE genes identified by the corresponding method. Genes with observed FDR smaller than $\alpha = 0.05$ were identified as DE. Both the CPLM and ZICP methods detect a comparable number of DE genes with scREhurdle. It is to be noted that, hurdle models due to their two-part formulation, also report genes that show significant difference in the proportion of zeros (i.e. based on the logistic sub-component), not explicitly tested by non-hurdle models including CPLM and ZICP. When the hypotheses are aligned for a fair comparison (i.e. testing for difference in mean expression using $H_0 : \beta = 0$ against $H_1 : \beta \neq 0$), this leads to a significant drop in both MAST- and scREhurdle-detected significant genes, as expected.

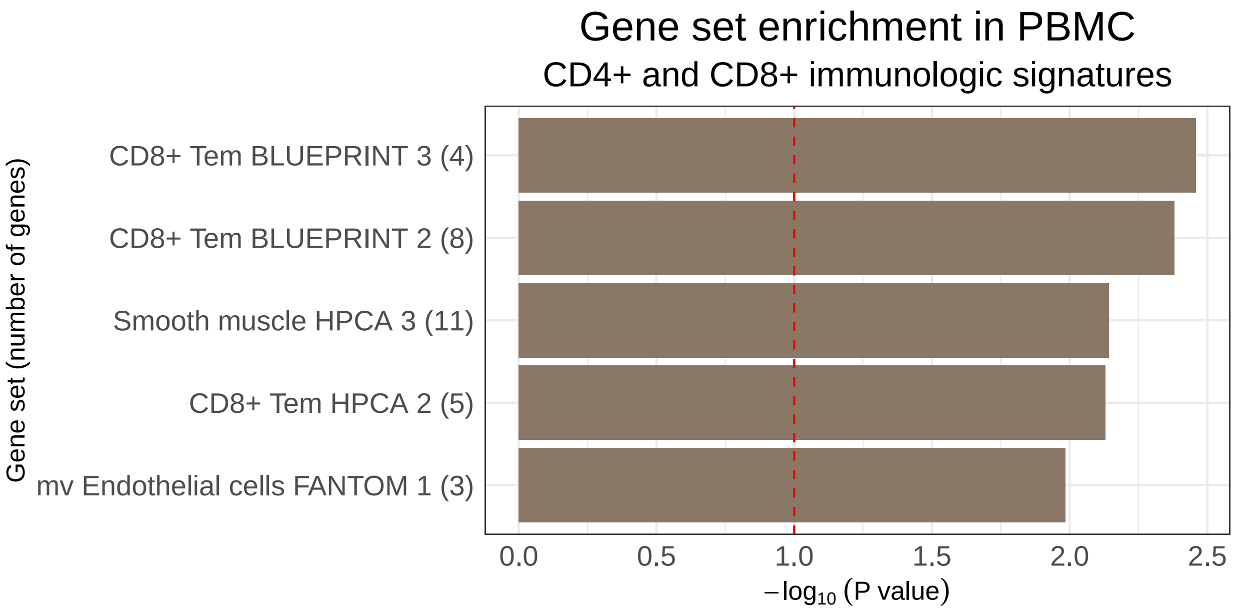


**Web Figure S11**: **Tweedieverse reveals biologically meaningful gene sets that are closely related to CD4+ and CD8+ immune processes**. Statistically significant gene sets (immunologic signatures) were identified using Benjamini and Hochberg (1995) (observed FDR smaller than $\alpha = 0.10$) as identified by the permutation-based Kolmogorov–Smirnov (KS) test (based on 100,000 null permutations) on uniquely determined genes detected by Tweedieverse. The bars in the $x$-axis indicate the $-\log_{10}()$ of $p$-values calculated as the fraction of permutation values that are at least as extreme as the original KS statistic derived from the non-permuted data. Numbers in the parentheses indicate the size of the gene sets.
